# Late-life isoleucine restriction promotes physiological and molecular signatures of healthy aging

**DOI:** 10.1101/2023.02.06.527311

**Authors:** Chung-Yang Yeh, Lucas C.S. Chini, Jessica W. Davidson, Gonzalo G. Garcia, Meredith S. Gallagher, Isaac T. Freichels, Mariah F. Calubag, Allison C. Rodgers, Cara L. Green, Reji Babygirija, Michelle M. Sonsalla, Heidi H. Pak, Michaela Trautman, Timothy A. Hacker, Richard A Miller, Judith Simcox, Dudley W. Lamming

## Abstract

In defiance of the paradigm that calories from all sources are equivalent, we and others have shown that dietary protein is a dominant regulator of healthy aging. The restriction of protein or the branched-chain amino acid isoleucine promotes healthspan and extends lifespan when initiated in young or adult mice. However, many interventions are less efficacious or even deleterious when initiated in aged animals. Here, we investigate the physiological, metabolic, and molecular consequences of consuming a diet with a 67% reduction of all amino acids (Low AA), or of isoleucine alone (Low Ile), in male and female C57BL/6J.Nia mice starting at 20 months of age. We find that both diet regimens effectively reduce adiposity and improve glucose tolerance, which were benefits that were not mediated by reduced calorie intake. Both diets improve specific aspects of frailty, slow multiple molecular indicators of aging rate, and rejuvenate the aging heart and liver at the molecular level. These results demonstrate that Low AA and Low Ile diets can drive youthful physiological and molecular signatures, and support the possibility that these dietary interventions could help to promote healthy aging in older adults.

## Introduction

The development of effective geroprotective therapies is necessary to prevent and delay diseases of aging in the rapidly graying population of our era. One of the most robust methods of promoting health and lifespan in diverse species is calorie restriction (CR), in which animals’ *ad libitum* caloric intake is restricted by 20-40% (Green et al., 2022a). While CR is highly effective in abating many age-related diseases in animal models, long-term adherence to a reduced-calorie diet without malnutrition can be challenging (Mihaylova et al., 2023). Further, CR is less beneficial when initiated later in life (Hahn et al., 2019), which may limit its usefulness for older adults.

Contrary to the conventional wisdom that calories from different sources are equivalent, several retrospective and prospective clinical trials have found that eating a diet with lower levels of protein is associated with lower rates of age-related diseases, including cancer and diabetes, as well as with an overall reduction of mortality in those under age 55 (Levine et al., 2014; Sluijs et al., 2010). While the effect of long-term protein restriction (PR) on human aging has not been tested in a randomized clinical trial (RCT), short-term RCTs in people who are overweight or have diabetes have found PR to promote metabolic health, reducing adiposity and improving glycemic control (Ferraz-Bannitz et al., 2022; Fontana et al., 2016). PR has also been repeatedly shown to increase the health and lifespan of model organisms, including flies and rodents (Hill et al., 2022; Mair et al., 2005; Richardson et al., 2021; Solon-Biet et al., 2014; Solon-Biet et al., 2015).

While the mechanism by which PR promotes healthy aging remains elusive, PR necessarily reduces dietary levels of the nine essential amino acids (EAAs). The EAAs are critical regulators of the metabolic response to PR in mice, and restriction of the nine EAAs is required for the benefits of a CR diet on lifespan (Yoshida et al., 2018). We and others have shown that restricting dietary levels of the three branched-chain amino acids (BCAAs; leucine, isoleucine, and valine) improves metabolic health in mice and rats, improving glucose tolerance and reducing adiposity in both lean and obese animals (Cummings et al., 2018; Richardson *et al*., 2021; White et al., 2016; Yu et al., 2021). The restriction of BCAAs extends lifespan and reduces frailty in male mice, while dietary supplementation with additional BCAAs shortens lifespan in both male and female mice (Richardson *et al*., 2021; Solon-Biet et al., 2019).

While most studies have investigated the physiological role of the BCAAs as a group, it is now apparent that the individual BCAAs have distinct physiological, metabolic, and molecular roles. In particular, we have shown that reducing dietary levels of isoleucine is both necessary and sufficient for many of the beneficial effects of PR in young mice, including improved glucose tolerance and reduced adiposity, and augments metabolism by increasing both food consumption and energy expenditure (Yu *et al*., 2021). In humans, dietary isoleucine levels, but not levels of leucine or valine, are strongly correlated with BMI (Yu *et al*., 2021). Finally, restriction of isoleucine in adult mice extends lifespan in both males and females (Green et al., 2023), and in humans, blood levels of isoleucine, but not of leucine or valine, are positively associated with mortality (Deelen et al., 2019). Thus, there is significant evidence that dietary isoleucine is a critical regulator of metabolic health and aging in both mice and humans.

Previous studies of isoleucine or protein restriction have primarily been conducted on mice treated with these diets at or prior to 6 months of age; in other words, in young mice. However, many geroprotective interventions like CR have reduced benefits when starting in mid-life or later (Hahn *et al*., 2019). To better understand the effects of restricting protein or isoleucine late in life, we placed 20-month-old C57BL/6J.Nia mice of both sexes on diets in which either all amino acids (Low AA) or isoleucine alone (Low Ile) was restricted by 67%. At this age, C57BL/6J mice are estimated to be roughly equivalent to 60-year-old humans (Flurkey et al., 2007). We tracked the weight, body composition, fitness, and frailty of these mice longitudinally over four months, examining the effects of the diets on physiology, glycemic control, and energy balance. We found that in both sexes, Low AA and Low Ile diets initiated late in life improve metabolic health and increase energy expenditure, with mixed effects on frailty and fitness. We found that both Low AA and Low Ile diets favorably affect a set of molecular aging rate indicators known to be altered by other lifespan-extending therapies. Finally, we found that a Low Ile diet favorably remodels the aging heart, with alterations in phosphatidylglycerol lipids, and reverses most age-associated changes in hepatic gene expression in both males and females. Our findings suggest that restricting dietary protein or isoleucine may promote healthy aging even when these diets started later in life.

## Results

### Restriction of isoleucine or all amino acids reduces body weight and adiposity in aged mice

To examine the effects of protein restriction (PR) and isoleucine restriction as late-life interventions, we obtained 20-month-old male and female C57BL/6J.Nia mice from the National Institute on Aging (NIA) Aged Mouse Colony, re-housed animals in larger groups to 2-3 animals per cage, and then randomized the mice of each sex to one of three experimental groups of equivalent body weight, adiposity, and frailty. Each group was placed on an amino acid (AA) defined diet containing all twenty common AAs; the diet composition of the Control diet (Control; TD.140711) reflects that of a chow diet in which 22% of calories are derived from protein. The other two groups were placed on diets in which either isoleucine was specifically reduced by 67% (Low Ile; TD.160734), or in which all twenty amino acids were reduced by 67% (Low AA; TD.140712). All three diets are isocaloric, with identical levels of fat; in the Low Ile diet, non-essential AAs were increased to keep the calories derived from AAs constant, while in the Low AA diet, carbohydrate levels were increased. All three of these diets have been used previously (Green *et al*., 2023; Yu *et al*., 2021), and their detailed composition can be found in **Table 1**. We placed an additional group of 6-month-old mice from the NIA Aged Mouse Colony on the Control diet as a young control group (Young Control).

**Table 1.**
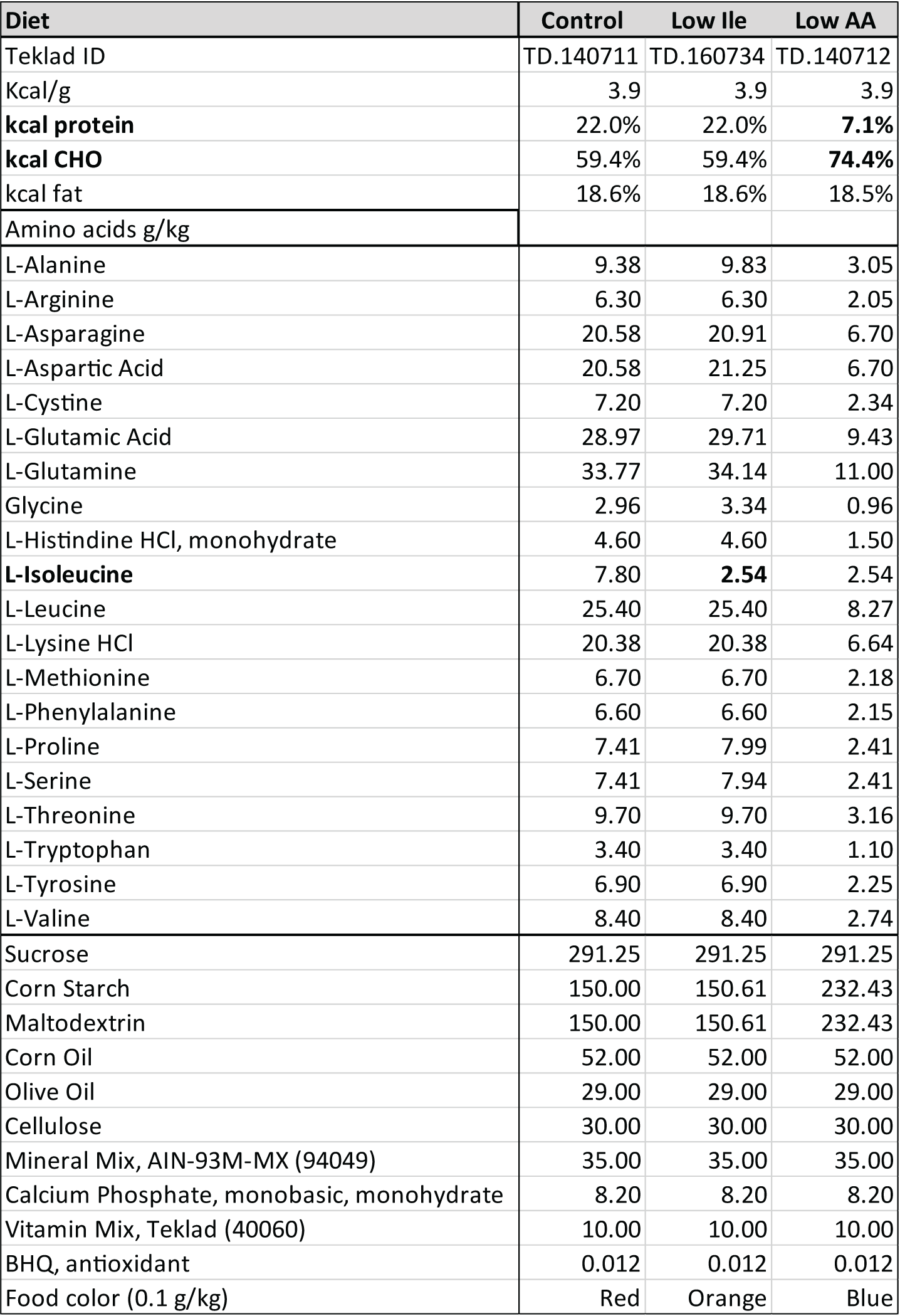
Diet composition. Diet composition and calorie content for diets used in this study.

We monitored the mice longitudinally for 4 months with metabolic and behavioral phenotyping and periodic assessment of weight, body composition, frailty, and food intake (**Fig. 1A**). Aged Control-fed mice maintained their weight throughout the course of the experiments, while Aged Low Ile-fed and Aged Low AA-fed males had a significant decrease in body weight (**Fig. 1B**). Aged Low Ile-fed males experienced rapid weight loss during the first 2-3 weeks before stabilizing, while the weight of Aged Low AA-fed males declined more gradually (**Fig. 1B**). The weight loss of Aged Low Ile-fed males was the result of decreases in both fat and lean mass, whereas the weight loss of Aged Low AA-fed males was the result only of lean mass loss (**Figs. 1C-E**). Both Low Ile and Low AA-fed males had an overall decrease in adiposity at every time point examined (**Fig. 1F**). The changes in weight and body composition were not due to decreased food consumption; at all times, the weight-normalized food consumption of Aged Low Ile- and Aged Low AA-fed mice was equal to or greater than Aged Control-fed males (**Fig. 1G**).

**Figure 1.**
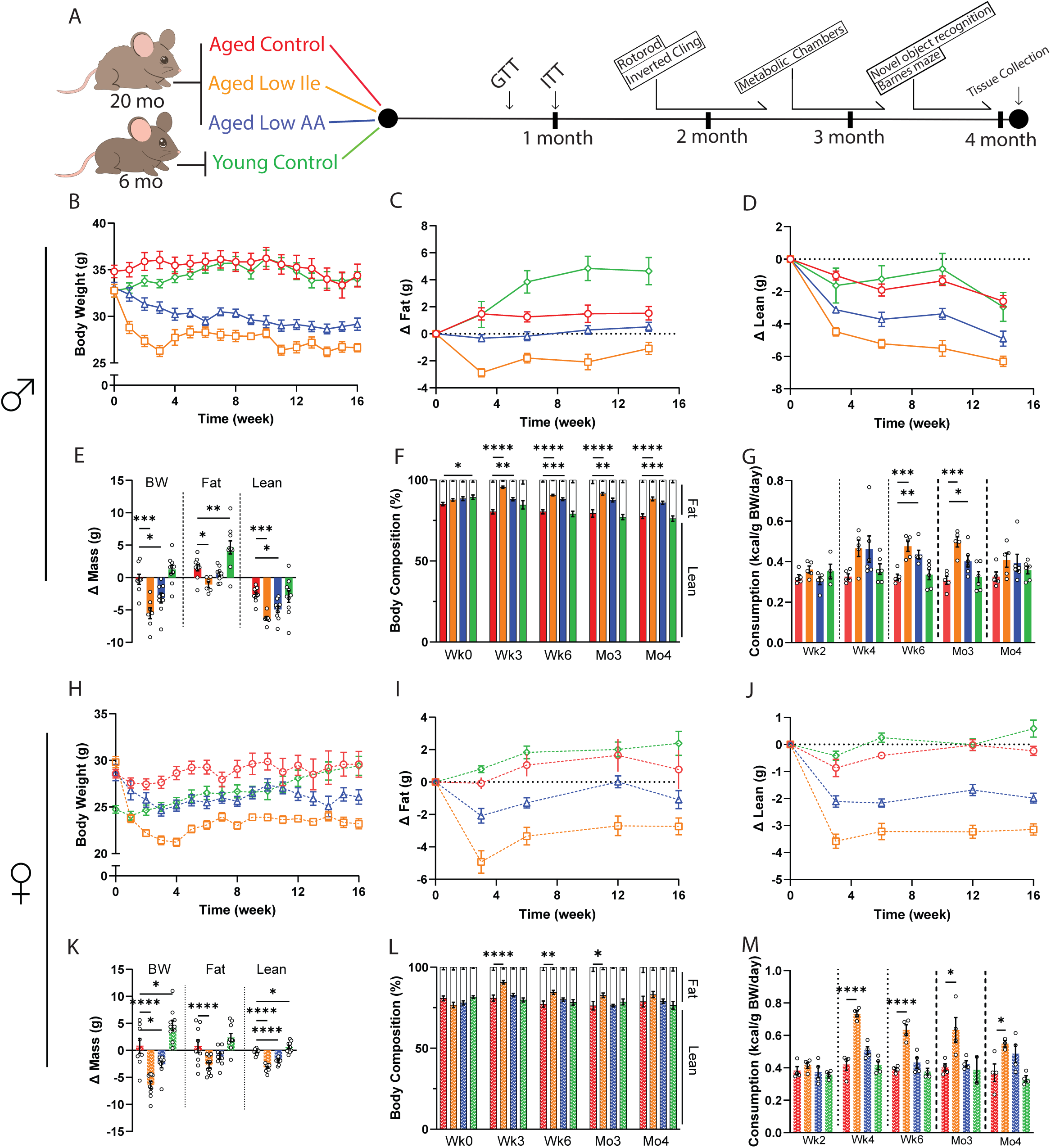
Low Ile and Low AA diets promote leanness in aged C57BL/6J.Nia mice. **(A)** Experimental scheme. Three different amino acid defined diets were utilized: Control, Low Ile, and Low AA. Aged mice began their respective diets at 20 months of age, while Young mice were fed the Control diet starting at 6 months of age. **(B-F)** Body weight (B), with change in fat mass (C) and lean mass (D) of male mice was tracked over time. (E) Change in body weight, fat, and lean mass during the course of the experiment. (F) Body composition percentage. (B-F) n=10-13/group; (E-F) ANOVA followed by Dunnett’s test vs. Aged Control-fed mice. **(G)** Food consumption of male mice throughout the experiment (n=5-6 cages/group, ANOVA followed by Dunnett’s test vs. Aged Control-fed mice. **(H-L)** Body weight (H), with change in fat mass (I) and lean mass (D) of female mice was tracked over time. (J) Change in body weight, fat, and lean mass during the course of the experiment. (K) Body composition percentage. (H-L) n=10-11/group; (K-L) ANOVA followed by Dunnett’s test vs. Aged Control-fed mice. **(M)** Food consumption of female mice throughout the experiment. n=2-4 cages/group, ANOVA followed by Dunnett’s test vs. Aged Control-fed mice. *p<0.05, **p<0.01, ***p<0.001, ****p<0.0001. Data presented as mean ± SEM.

We observed similar effects of Low Ile and Low AA diets on the weight and body composition of aged female mice (**Figs. 1H-K**). The overall effect of the Low Ile diet in females was similar to what we observed in males, with an overall reduction in adiposity, while in females the effect of a Low AA diet on adiposity was not significant (**Fig. 1L**). As in males, the weight loss of Low Ile and Low AA-fed female mice was not the result of reduced food intake, as the caloric intake of these groups was at all times equal to or greater than Aged Control-fed females (**Fig. 1M**).

### Late-life restriction of isoleucine or all amino acids improves aspects of healthspan

Decreased protein intake is associated with frailty and sarcopenia in older adults (Coelho-Junior et al., 2020; Coelho-Junior et al., 2018); however, we have found that long-term restriction of all amino acid, the BCAAs, or isoleucine alone actually reduces overall frailty in mice (Green *et al*., 2023; Richardson *et al*., 2021). To assess the impact of beginning a Low AA or Low Ile diet later in life, we utilized a validated mouse frailty index that quantifies frailty through the accumulation of deficits (Whitehead et al., 2014).

We initially performed a 3-way mixed-effects analysis to identify effects of Age, Diet, and Sex for each group of mice as compared to the Aged Control animals. As sex was a significant factor in the response of Low Ile-fed mice (**Sup. Figs. 1A-C**), we analyzed the data in a pairwise 2-way mixed-effects analysis with the sexes separated. As expected, we observed increased frailty in Aged Control-fed mice relative to Young Control-fed mice in both males and females, and we observed that frailty increased in Aged Control-fed mice as they grew older (**Figs. 2A, H**). While we did not observe an overall significant difference in frailty in either sex of mice fed a Low Ile diet, there was a trend of Low Ile-fed males towards reduced frailty (p=0.071). Similar non-significant trends were observed in the frailty of Aged Low AA-fed males and females (p=0.11 and p=0.18, respectively). The categorical parameters and the individual measures that contributed the most to these group trends were analyzed, and we found that Low AA and Low Ile diets specifically reduced deficits in body condition and distended abdomen in males but not females (**Figs. 2B-C, I-J**, **Sup. Figs. 1D-V**).

**Figure 2.**
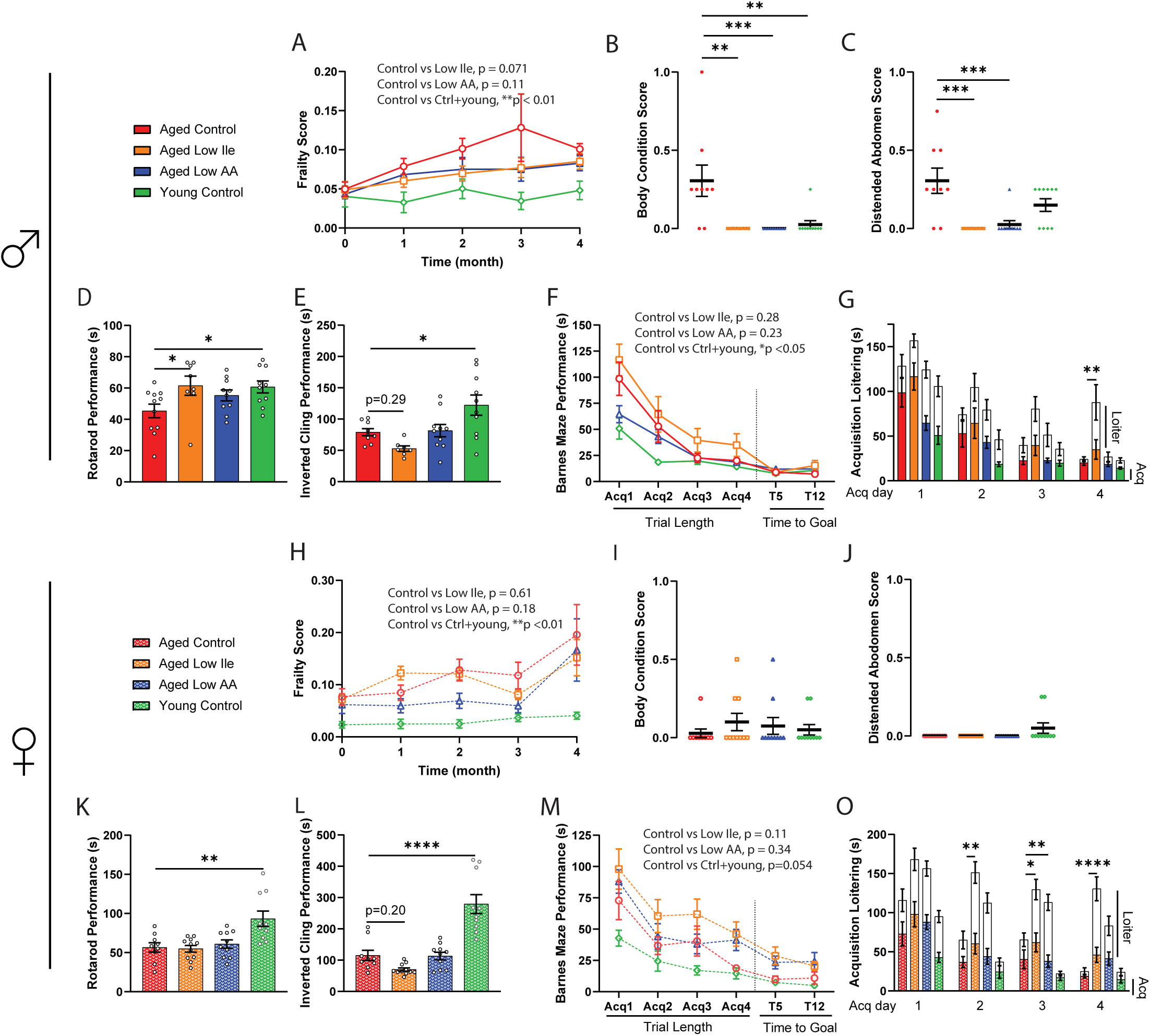
Late-life feeding of Low Ile and Low AA diets promote aspects of healthspan, particularly in male mice. **(A-C)** Frailty score of male mice (A) was tracked throughout the experiment between 20 and 24 months of age (n=10-13/group, p-value represents the result of the indicated 2-way mixed-effects analysis). (B-C) Selected individual frailty categories, presented as the average of scores during the 3^rd^ and 4^th^ month of the experiment. (A-C) n=10-13/group at the beginning of the experiments, ANOVA followed by Dunnett’s test vs. Aged Control-fed mice. **(D-E)** Male rotarod (D) and inverted cling (E) performance were assessed between 22-23 months of age (n=8-11/group, ANOVA followed by Dunnett’s test vs. Aged Control-fed mice). **(F-G)** Male Barnes Maze Test performance at 24 months of age (F). n=7-10/ group, p-value represents the effect of diet in the indicated 2-way ANOVA, acquisition time only. (G) Barnes Maze Test acquisition trial duration with loitering (test on loitering time only, ANOVA followed by Dunnett’s test vs. Aged Control-fed mice). **(H-J)** Frailty score of female mice (H) was tracked throughout the experiment between 20 and 24 months of age (n=10-11/group, p-value represents the effect of diet in the indicated 2-way ANOVA). (I-J) Selected individual frailty categories, presented as the average of scores during the 3^rd^ and 4^th^ month of the experiment. (H-J) n=10-11/group at the beginning of the experiments, ANOVA followed by Dunnett’s test vs. Aged Control-fed mice. **(K-L)** Female rotarod (K) and inverted cling (L) performance were assessed between 22-23 months of age (n=8-11/group, ANOVA followed by Dunnett’s test vs. Aged Control-fed mice). **(M-O)** Female Barnes Maze Test performance at 24 months of age (M). n=8-10/ group, p-value represents the effect of diet in the indicated 2-way ANOVA, acquisition time only. (O) Barnes Maze Test acquisition trial duration with loitering (test on loitering time only, ANOVA followed by Dunnett’s test vs. Aged Control-fed mice). *p<0.05, **p<0.01, ***p<0.001, ****p<0.0001. Data presented as mean ± SEM.

We assessed neuromuscular coordination and muscle strength with an accelerating rotarod and an inverted cling test. As expected, Young Control-fed males and females performed better on these assays than Aged Control-fed mice (**Figs. 2D-E, K-L**). Aged Low Ile-fed males performed significantly better in the rotarod test than Aged Control-fed males, and the performance of both Aged Low Ile-fed and Aged Low AA-fed males was comparable to the Young Control-fed males (**Fig. 2D**). In contrast, there was no effect of diet on the rotarod performance of aged females (**Fig. 2K**). In an inverted cling test, there was a non-significant trend towards decreased cling time in Aged Low Ile-fed male and female mice (p=0.29 and p=0.20 respectively), and no change in Low AA-fed mice of either sex (**Figs. 2E and L**). As Aged Low Ile-fed mice are lighter than Aged Control-fed mice, we analyzed rotarod and cling test performance with body weight as a covariate (**Sup. Figs. 2A-D**); we found that the inverted cling performance of Aged Low Ile-fed females was significantly worse compared to Aged Control-fed females (**Sup. Fig. 2D**). To evaluate whether this trend of diet-induced loss in grip strength is age-dependent, a separate cohort of young 3-month-old male mice was fed the Low Ile diet for 2 months. We observed a similar trend (p=0.053) of reduced cling performance in Low Ile-fed males despite their young age and decreased body weight (**Sup. Figs. 2E-F**).

C57BL/6J mice suffer from age-related cognitive decline (Majumder et al., 2012). We examined the impact of Low Ile and Low AA diets on memory by conducting a Novel Object Recognition test (NOR) and a Barnes Maze Test (BMT) at approximately 23 months of age. During the habituation phase for the NOR, we also quantified the spontaneous movements of each group in the open field. Interestingly, Aged Control-fed females exhibited increased locomotion compared to both Aged Low Ile-fed females and the Young Control-fed females; this effect of diet and age was not observed in male mice (**Sup. Figs. 3A, B**). In the NOR test, we did not observe a significant effect of age on performance during the acquisition phase, the short-term memory (STM) test, or the long-term memory (LTM) test between Young Control-fed and Aged Control-fed mice of either sex, and thus it is not surprising that we also did not observe significant diet-induced differences (**Sup Figs. 3C-E, G-I**). Summing all three phases, the total time spent investigating both objects was the highest in Young Control-fed males and significantly lower in Aged Control-fed males, with Aged Low Ile-fed males investigating the objects for even less time (**Sup** **Fig. 3F**). There was no group effect in the total investigation time of the females (**Sup** **Fig. 3J**).

**Figure 3.**
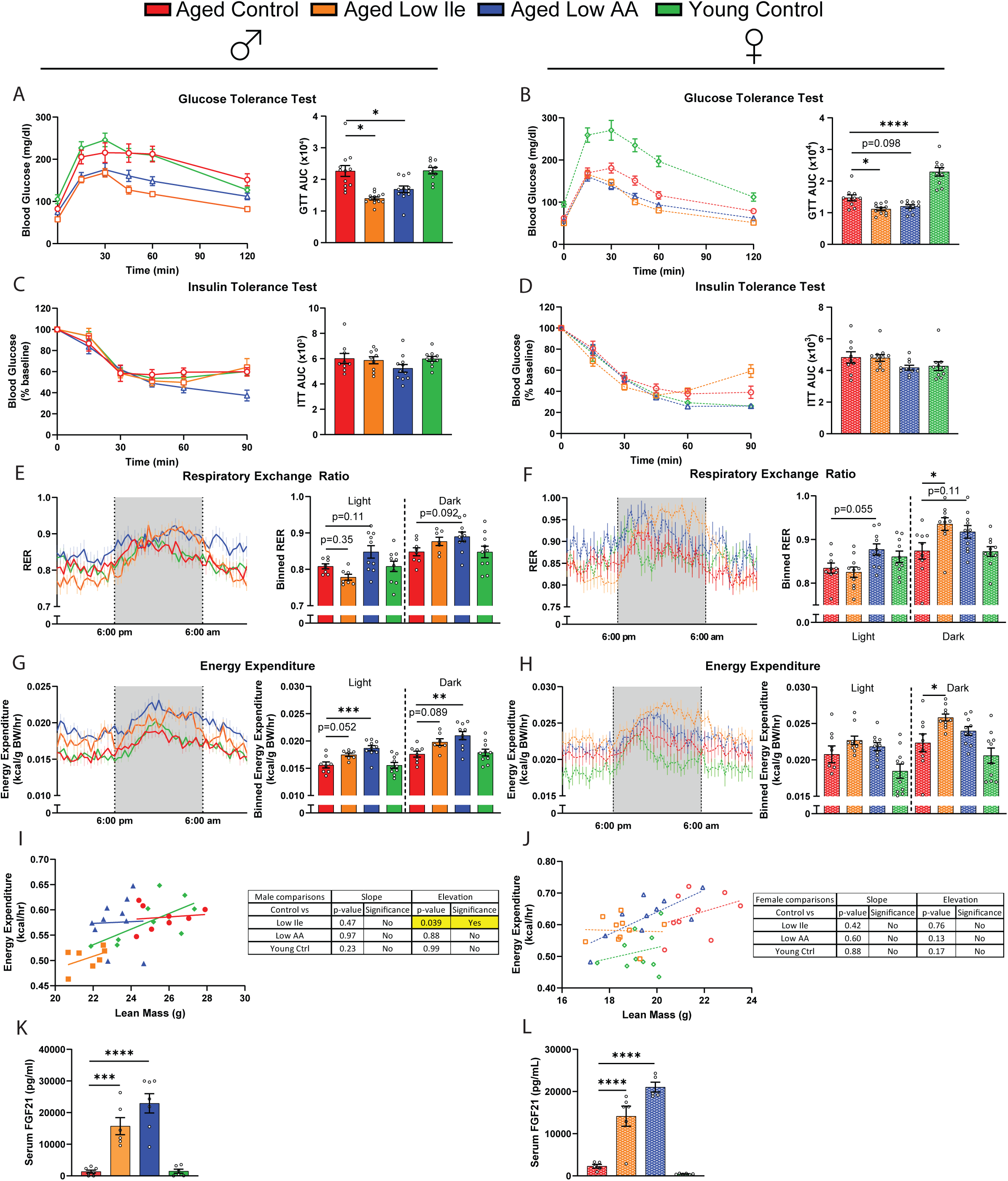
Late-life feeding of a Low Ile or Low AA diet improves glycemic control and boosts energy expenditure. **(A-D)** Glucose tolerance test in male (A) and female (B) mice fed the indicated diets. Insulin tolerance test in male (C) and female (D) mice fed the indicated diets. n=10-13/group, ANOVA followed by Dunnett’s test vs. Aged Control-fed mice. **(E-H)** Metabolic chambers were used to determine the respiratory exchange ratio (RER) male (E) and females (F) and the energy expenditure normalized to body weight in males (G) and females (H). n=7-10/group, ANOVA conducted separately for the light and dark cycles followed by Dunnett’s vs. Aged Control-fed mice. **(I-J)** ANCOVA of energy expenditure with lean mass as a covariate in males (I) and females (J). n=7-10/group. **(K-L)** The serum FGF21 level at the end of the experiment, after 16 hr fasting overnight and 3 hr refeeding. n=5-7/group, ANOVA followed by Dunnett’s test. *p<0.05, **p<0.01, ***p<0.001, ****p<0.0001. Data presented as mean ± SEM.

In the BMT, we observed an age-related deficit during the acquisition phase in both sexes of mice (**Figs. 2F, M**). Aged Low Ile-fed mice of both sexes trended towards performing more poorly than Aged Control-fed mice (p=0.28 in males, p=0.11 in females). However, we noticed that many of the aged mice, particularly those fed the Low AA and Low Ile diets, could quickly locate the escape box, which ends the acquisition trial, but spent a significant amount of time hesitating instead of entering immediately. Manually measuring this additional period of loitering time, we found that loitering was persistently higher in Low Ile-fed males and females, and in in Low AA-fed females (**Figs. 2G, O**). In sharp contrast, Young and Aged Control-fed mice of both sexes became increasingly decisive and loitering decreased over the acquisition sessions (**Figs. 2G, O**).

### Restriction of isoleucine or all amino acids promote glucose tolerance and energy expenditure in aged mice

We have shown that the restriction of protein or isoleucine can promote glucose tolerance when these diets are initiated in young mice. Here, we examined the effect of these interventions on metabolic parameters when initiated in aged mice of both sexes. Aligned with our previous observations in young males, we found that consumption of either a Low AA or a Low Ile diet improved glucose tolerance regardless of sex (**Figs. 3A-B**). While conducting these assays, we observed that Aged Low Ile-fed males had significantly lower fasting blood glucose after 16 hr of fasting (**Sup. Figs. 4A-B**). In a separate cohort of 25-month-old animals, a Low Ile diet likewise significantly improved glucose tolerance in aged males and females (**Sup. Figs. 4C-D**). Consistent with our previous results in young males, neither a Low Ile diet nor a Low AA diet significantly improved insulin sensitivity as assessed via intraperitoneal administration of insulin (**Figs. 3C-D**).

As we observed significant changes to body weight and food intake, we utilized metabolic chambers to examine how energy balance was impacted by these diets. We have previously observed that restriction of all amino acids or isoleucine alone increases the respiratory exchange ratio (RER) and energy expenditure in young male mice (Yu *et al*., 2021), and we observed a similar trend towards increased RER in Aged Low AA-fed mice of both sexes (**Figs. 3E-F**). Aged Low Ile-fed mice of both sexes likewise had higher RER during the dark cycle, reaching statistical significance in the case of Aged Low Ile-fed females (**Fig. 3E-F**).

As we previously observed in young males, the energy expenditure of Aged Low Ile or Low AA-fed males was higher than that of Aged Control-fed males, with the Aged Low AA-fed mice having significantly increased energy expenditure during both the light and dark cycles (**Fig. 3G**). Energy expenditure was likewise significantly higher in Aged Low Ile-fed females during the dark cycle (**Fig. 3H**). Using ANCOVA analysis with lean mass as a covariate, we found that Aged Low Ile-fed males have reduced energy expenditure independent of their lean mass compared to Aged Control-fed males (**Figs. 3I-J**). This indicates that the differences in body composition and weight between the groups contribute greatly to the disparity seen in their energy expenditure. The differences in energy expenditure were not due to differences in spontaneous activity (**Sup. Figs. 4E-F**), but were accompanied by induction of the energy balance hormone FGF21 (**Figs. 3K-L**).

### Restriction of isoleucine or all amino acids selectively ameliorate molecular indicators of aging rate

Recently, a panel of molecular indicators of aging rate that is remarkably conserved across multiple lifespan-extending treatments has been identified (Miller et al., 2023). To determine the effect of late-life Low Ile and Low AA on these pathways, we generated liver protein lysates from mice that had been on these or Control diet at 24 months of age, after consuming the diets for 4 months, and immunoblotted using antibodies against these aging rate indicators, which were then quantified (**Fig. 4A-I**).

**Figure 4.**
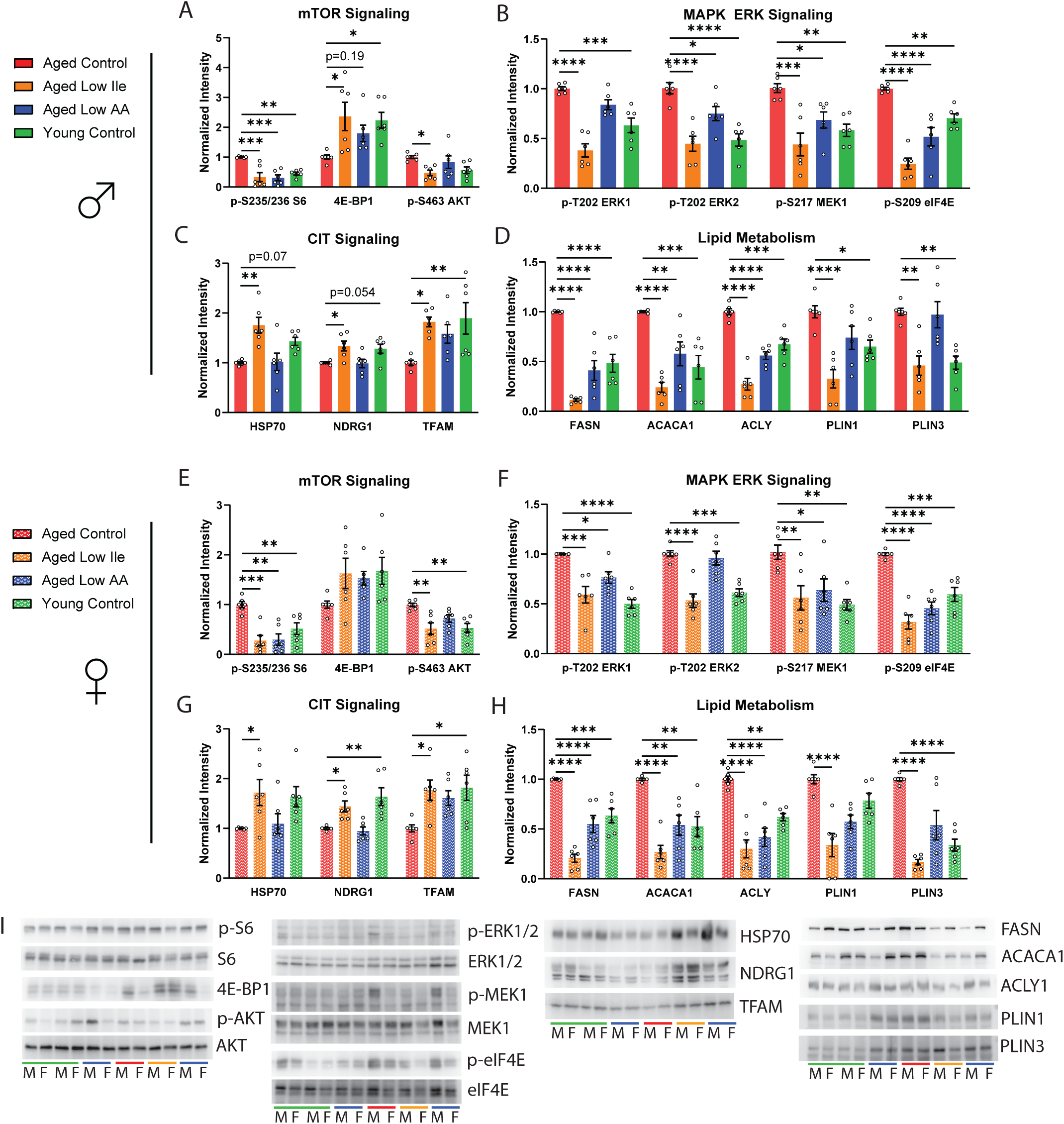
Low Ile and Low AA diet ameliorates multiple molecular indicators of aging rate in the liver. **(A-D)** Diet and age alters aging rate indicators related to (A) mTOR signaling, (B) MAPK ERK signaling, (C) cap-independent translation (CIT), and (D) lipid oxidation in the liver of male mice. **(E-H)** Diet and age alters aging rate indicators related to (E) mTOR signaling, (F) MAPK ERK signaling, (G) cap-independent translation (CIT), and (H) lipid oxidation in the liver of female mice. **(I)** Representative Western blots of the proteins analyzed. n=6/group, ANOVA followed by Dunnet’s test. *p<0.05, **p<0.01, ***p<0.001, ****p<0.0001. Data presented as mean ± SEM.

mTORC1 is a master regulator of growth and nutrient sensing, and is activated by BCAAs (Simcox and Lamming, 2022). mTORC1 activity increases with age and its pharmacological inhibitor rapamycin is a robust geroprotector (Baar et al., 2016; Mannick and Lamming, 2023). Here, we found that Aged Control-fed males exhibit significantly higher levels of phosphorylated S6, a downstream readout of mTORC1 activity, than Young Control-fed males; this age-associated increase was suppressed in both Aged Low Ile- and Low AA-fed males (**Fig. 4A**). The mTORC1-controlled translation regulator 4E-BP1 did not have significant changes in its phosphorylation ratio but total 4E-BP1 expression was significantly decreased by aging and this was significantly rescued by a Low Ile diet and improved by Low AA diet (p=0.19, **Fig. 4A**; **Sup. Fig 5A**). The same effects of age and dietary interventions on these mTORC1 targets were also observed in the females with the effect size being slightly less robust with 4E-BP1 (**Fig. 4E**). Only the Low Ile diet suppressed the age-dependent increase in the phosphorylation of the mTORC2 target AKT S473 (**Fig. 4A, E**). We did not observe significant changes in the phosphorylation of ULK1 S757 and eIF2α S51 across all groups (**Sup. Fig. 5A-B**). In females only, the phosphorylation of the mTORC1 substrate S6K1 exhibited a strong trend of being suppressed by a Low AA diet (p=0.078, **Sup. Fig 5B**).

The MAPK ERK signaling pathway regulates translation, increases with age, and can be suppressed by the lifespan-extending treatments canagliflozin and 17aE2 (Jiang et al., 2023; Miller et al., 2020). We found that Aged Control-fed male mice have significantly increased phosphorylation of ERK1 T202, ERK2 T202, MEK1 S217, as well as eIF4E S209 compared to the Young Control-fed mice (**Fig. 4B**), similar to previous observations in UM-HET3 mice. The Low Ile diet robustly decreased phosphorylation of all four of these substrates, while the Low AA diet suppressed all but ERK1 T202 phosphorylation.

An increase in cap-independent translation (CIT) is a shared feature in long-lived genetic and drug intervention models (Shen et al., 2021). We observed Aged Control-fed male mice has a non-significant decrease of HSP70 and NDRG1 (p=0.07 and p=0.054 respectively), and a significant decrease in TFAM (**Fig. 4C**). Remarkably, all three of these proteins were significantly increased in the Aged Low Ile-fed animals, but not in the Aged Low AA-fed animals. Recent transcriptomic analysis has revealed that systemic shifts in fatty acid oxidation, particularly in the liver, are associated with lifespan-extending interventions (Watanabe et al., 2023). We identified age-dependent increases in the fatty acid regulators FASN, ACACA1, ACLY, PLIN1, and PLIN3 (**Fig. 4D**). A Low Ile diet in Aged male mice decreased the expression of all five proteins while a Low AA diet decreased expression of FASN, ACACA1, and ACLY.

In the female liver, Low Ile and Low AA diets had similar molecular effects as in males with respect to MAPK ERK signaling, CIT signaling, and lipid metabolism molecular pathways (**Fig. 4F-H**). Overall, both Low Ile and Low AA diets appear to share many, but not all, molecular signatures with each other and with other life-extending interventions.

### Restriction of isoleucine induces sex-specific cardiac remodeling

Mice as well as humans undergo age-associated cardiac hypertrophy and diastolic dysfunction, which can be reversed or ameliorated by calorie restriction as well as by inhibition of mTORC1 signaling (Dai et al., 2014). To determine if a Low Ile or a Low AA diet affect cardiac function, we performed echocardiography to evaluate heart function in a separate cohort of animals at 25 months of age after 6 weeks of dietary intervention.

We identified a number of sex-specific and diet-induced changes to cardiac parameters (**Fig. 5**, **Table 2**). In aged males, a Low Ile diet increased left ventricle (LV) mass while decreasing mean and peak aortic flow velocity relative to Aged Control-fed males (**Table 2**). These changes were absent in Aged Low Ile-fed females (**Table 2**). In contrast, we detected an age-dependent increase in the LV posterior wall thickness and the stroke volume of Aged Control-fed females compared to the Young Control-fed females (**Fig. 5A, C**). Aged Low Ile-fed females had a significant decrease in diastolic LV inner diameter, decreased stroke volume, and increased heart rate relative to Aged Control-fed females (**Figs. 5B-D**). In contrast, the Aged Low AA-fed female mice only had increased heart rate compared to the Aged Control-fed females (**Figs. 5D**). There were no significant changes in cardiac output in either sex (**Table 2**, **Fig. 5E**). Overall, dietary intervention caused clear sex-specific changes in the cardiac parameters in aged mice, but these effects induced by Low Ile in Aged females clearly trend towards the level of the Young Control-fed mice (**Fig. 5B-E**).

**Figure 5.**
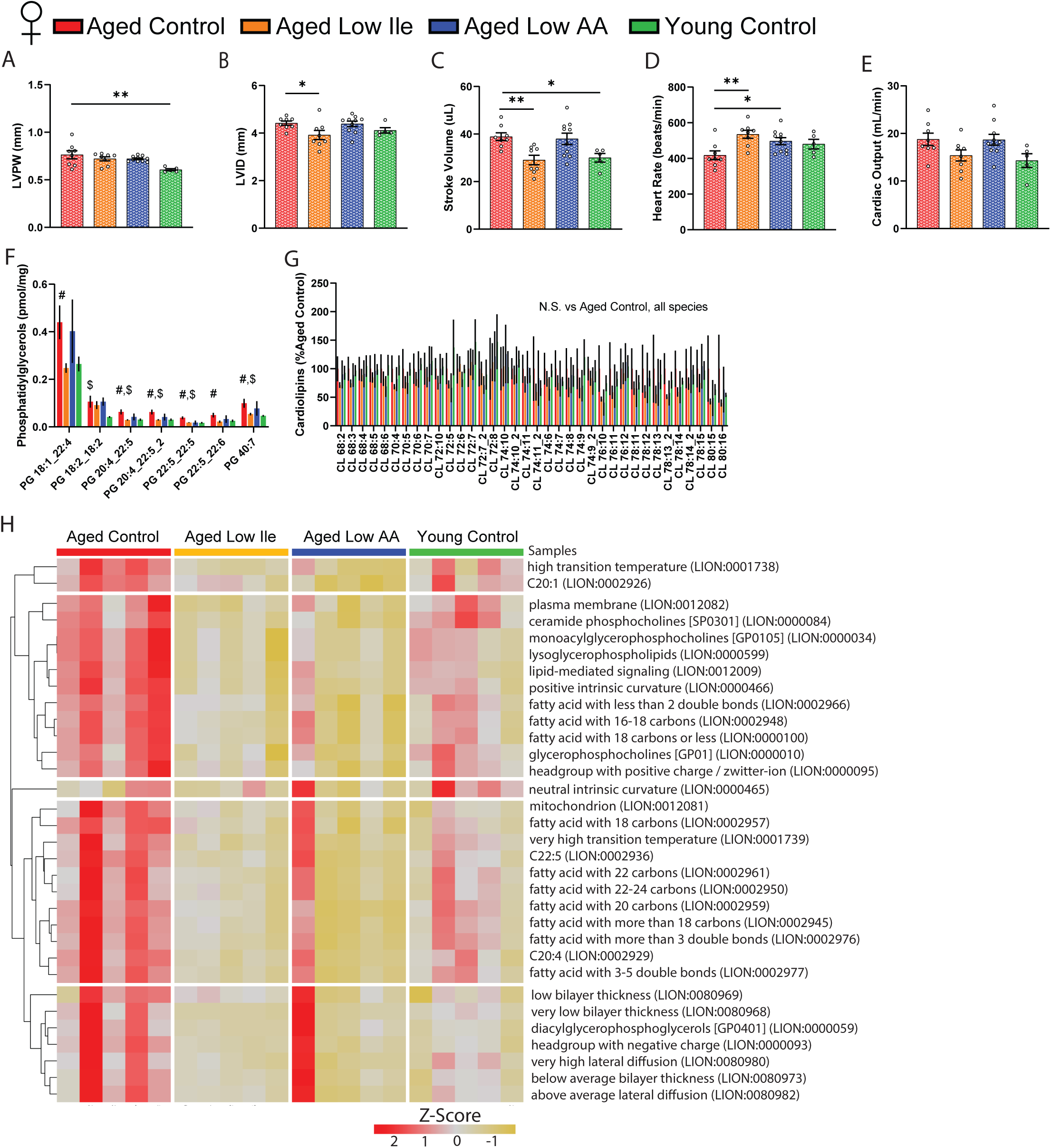
Low Ile diet promotes youthful functional and molecular aspects of the female mice heart. **(A-E)** Echocardiogram evaluation of female mice at 25 months of age. (A) Left ventricle posterior wall diameter, (B) left ventricle inner diameter, (C) stroke volume, (D) heart rate, and (E) cardiac output. n=5-10/group, *p<0.05, **p<0.01, ANOVA followed by Dunnett’s test. **(F-G)** Statistically significant phosphatidylglycerols (F) and all cardiolipins (G) in female mice hearts at 24 months of age after 4 months of dietary intervention. n=5/group, #p<0.05 Aged Control vs. Aged Low Ile, $p<0.05 Aged Control vs. Young Control, t-test. **(H)** LION lipid ontology analysis of significantly altered lipid species in the female mice heart. Data presented as mean ± SEM.

**Table 2.**
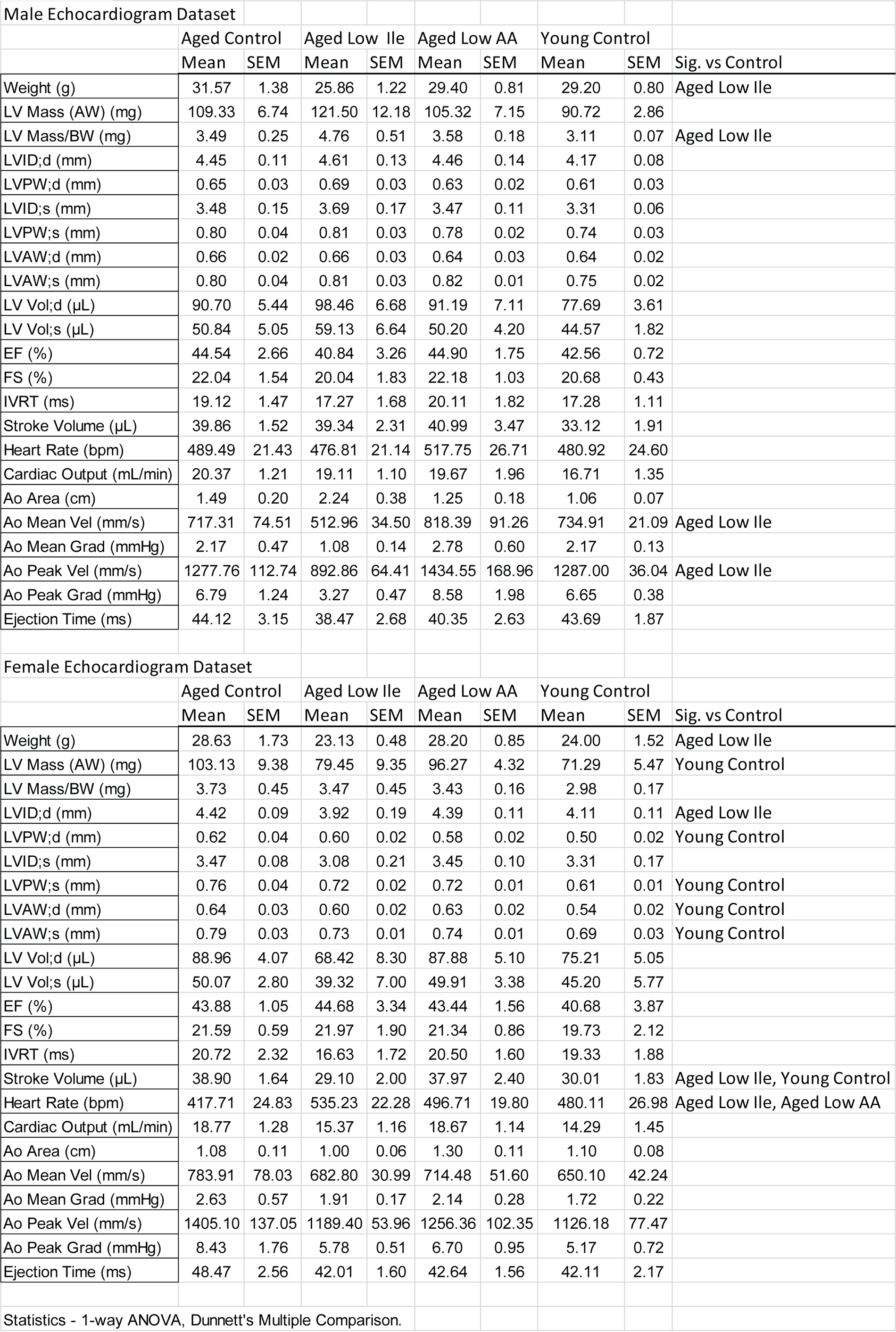
Echocardiography results. Detailed male and female echocardiogram dataset in mice after 6 weeks of dietary intervention starting from 24 months of age.

To obtain an unbiased and comprehensive assessment of the effects of Low Ile and Low AA diets on the aged female heart, we performed lipidomic profiling on the hearts of female mice collected at 24 months of age following 4 months of dietary intervention. Lipids are necessary components of cellular membranes, and the lipid composition is distinct for cell types and different organelles. We observed a significant age-dependent increase in several phosphatidylglycerol species that was mostly ameliorated in Low Ile-fed mice, but not in Low AA-fed mice (**Fig. 5F**). As phosphatidylglycerol is the primary component of the mitochondrial outer membrane, we also explored the most abundant mitochondrial inner membrane lipid and the phosphatidylglycerol-derivative, cardiolipin. Interestingly, all cardiolipin species were unchanged (**Fig. 5G**). The increase in phosphatidylglycerol without a change in cardiolipins could indicate a shift in mitochondrial morphology and would suggest increased fission which is associated with cardiomyopathy (Dorn, 2016). We performed lipid ontology enrichment analysis (LION) using all significantly altered lipid species; the enriched LION terms are shown in a heatmap (**Fig. 5H**). Overall, we observed a rejuvenating effect on the lipid profile of both Low Ile and Low AA-fed females, with most of the LION terms showing age-related increases and being restored to a more youthful level in Low Ile- and Low AA-fed mice. The list of all significant differentially expressed lipid species are provided in **Sup. Data Table**.

### Restriction of isoleucine or all amino acids induces senomorphic changes in the aged liver

Cellular senescence plays a large role in promoting several aging phenotypes of the liver, including fibrosis and non-alcoholic fatty liver disease (Matthew et al., 2017). To characterize the molecular remodeling that occurs in the aged liver induced by the two dietary interventions, we performed real time quantitative PCR (rt-qPCR) for gene expression level changes of well-established senescence-associated markers in male mice. Compared to the Aged Control-fed mice, Aged Low Ile-fed males had significantly increased expression of *p21* while Low AA-fed males had significantly reduced the expression of *Il-1a* and *Tnf-α* (**Sup. Fig. 6**). Of note, the trends in the changes of senescence markers induced by Low Ile and Low AA are distinct from each other, with Low Ile increasing and Low AA decreasing several canonical markers of senescence.

### Age-dependent differentially expressed genes in the liver are highly sensitive to a Low Ile diet

To obtain an unbiased and global view of the effects of isoleucine restriction, we performed transcriptional profiling of livers from Aged Control, Aged Low Ile, as well as Young Control mice of both sexes. We identified significantly differentially expressed genes (DEGs) induced by either old age and or by diet (**Fig. 6A-B**). In males, we identified 606 DEGs in response to aging (Aged Control vs Young Control) with 363 upregulated genes and 243 downregulated genes (**Fig. 6A**). In contrast, there were only 125 DEGs between Aged Low Ile-fed and Aged Control-fed males, with 54 upregulated genes and 71 downregulated genes (**Fig. 6B**). A similar effect was observed in females, with 911 DEGs upregulated and 676 downregulated in response to aging (**Fig. 6A**), and 2379 DEGs responded to Low Ile, with 1257 upregulated and 1122 downregulated genes (**Fig. 6B**).

**Figure 6.**
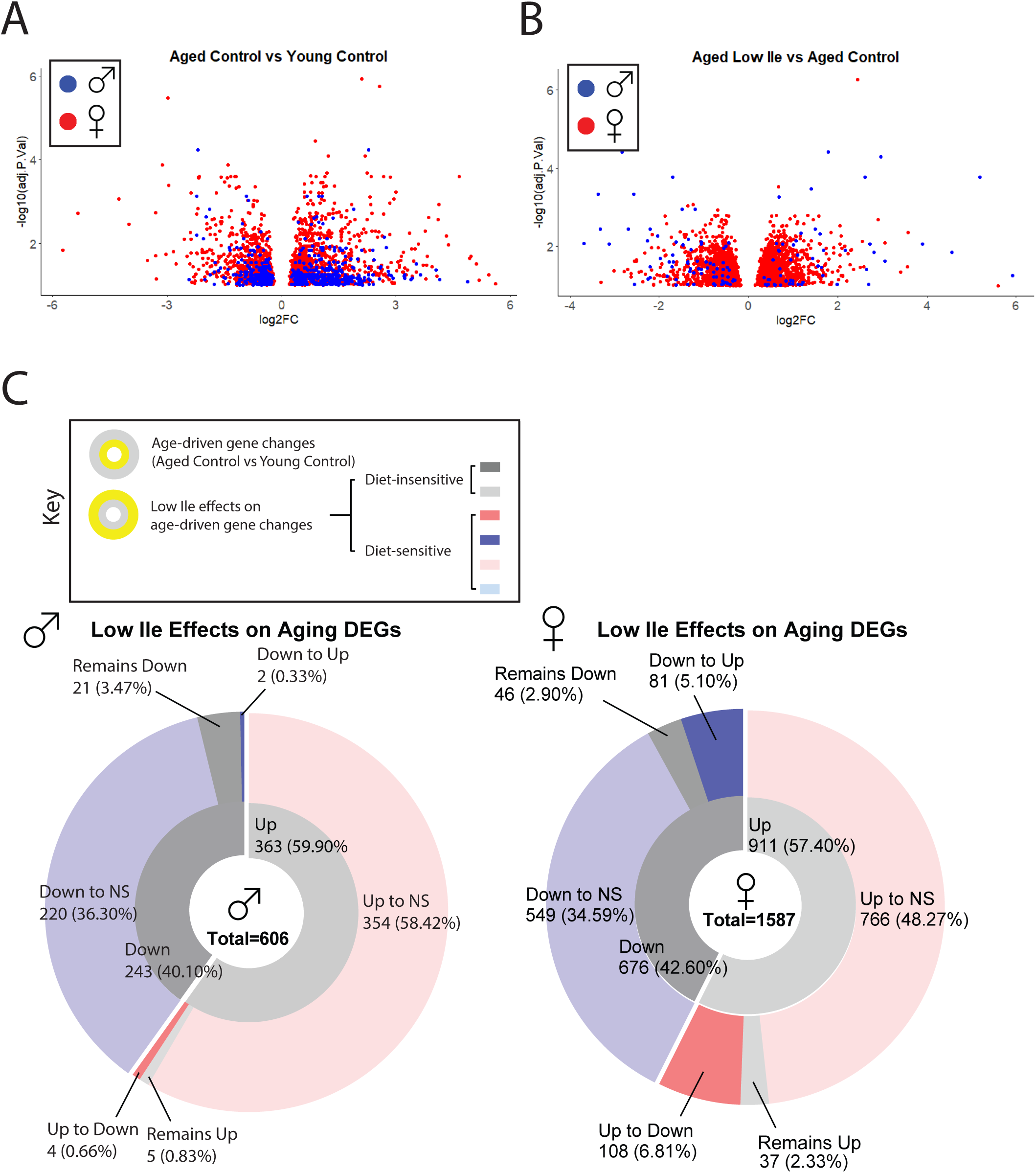
Late-life feeding of a Low Ile diet rejuvenates the liver transcriptome. **(A-B)** Volcano plots of differentially expressed genes in the liver of male (blue) and female (red) mice with age (A) and diet (B). All DEGs are determined at α = 0.10. **(C)** A summary of the effect of Low Ile on the age-driven differentially expressed gene sets in males (left) and females (right). Inner ring represents the genes up or downregulated by aging, while the outer ring represents the effect of a Low Ile diet on the DEGs altered with aging. n=5-6/group.

This pattern indicates that we should compare how the DEGs between Young Control-fed and Aged Control-fed mice differed from the DEGs between Young Control-fed and Aged Low Ile-fed mice. Of the 363 DEGs upregulated in Aged Control-fed males, 354 were not significantly different and 4 were significantly downregulated in Aged Low Ile-fed males. Of the 243 DEGs downregulated in Aged Control-fed males, 220 were not significantly differently expressed and 2 were significantly upregulated in Aged Low Ile-fed males (**Fig. 6C** **left**). A similar story was true in females, of the 911 significantly DEGs upregulated by age, the Low Ile diet caused 766 to become no longer significantly changed, and 108 reversed to downregulated. Of the 676 DEGs downregulated by age, Low Ile diet caused 549 to become no longer significantly changed, and 81 became upregulated. (**Fig. 6C** **right**). Overall, it seemed that a Low Ile diet “rejuvenates” the aging hepatic transcriptome in both sexes.

There were significantly more DEGs induced by both aging and diet in females than males, but most of the diet-induced DEGs in males presented a greater degree of change. The list of the top 50 DEGs induced by age and diet in both sexes are provided in **Sup. Data Table**. There was surprisingly little overlap between the male and female DEGs induced by a Low Ile diet (**Sup. Fig. 6A**). This resulted in drastically different pathway enrichment in our KEGG and GO analysis (**Sup. Fig. 6B-I**). While surprising, this agrees with previous research finding that there is a substantial effect of sex on the transcriptional changes induced by protein restriction or restriction of all three BCAAs (Green et al., 2022b; Richardson *et al*., 2021). A list of all significantly enriched KEGG pathway and GO terms are provided in **Sup. Data Table**.

## Discussion

Recent work from our lab and others has demonstrated that calorie quality, not just total quantity, is a critical determinant of biological health. Reducing dietary levels of protein, the three BCAAs, or isoleucine alone improves the metabolic health and extends the lifespan of mice (Fontana *et al*., 2016; Green *et al*., 2023; Richardson *et al*., 2021; Solon-Biet *et al*., 2019; Solon-Biet *et al*., 2014). However, all of these studies have initiated diets in relatively young mice; the ability of these interventions to promote healthy aging when begun in older animals is not clear. This is a critically important question, as to be of maximum clinical relevance, geroprotective interventions need to be able to promote healthy aging even when started late in life.

We and others have proposed the use of a comprehensive metabolic, physical, and cognitive phenotyping pipeline to assess the effects of interventions on the healthspan of aging mice (Bellantuono et al., 2020). Here, we have utilized this workflow to examine the effects of Low Ile and Low AA diets on healthspan when started in mice at 20 months of age, roughly equivalent to a 60-year-old human. Overall, we find that many effects of Low Ile and Low AA diets in aged mice are similar to those we observed in younger animals. In particular, Low Ile and Low AA diets have dramatic and largely positive effects on weight, body composition, glycemic control, and energy balance in aged mice of both sexes.

While there are significant metabolic benefits of Low Ile and Low AA diet in both sexes, there are also sex-specific effects on physical performance, frailty, and cognition that require further investigation. In contrast to our previous findings in young males and females, aged mice consuming either a Low Ile or Low AA diet have a rapid and significant loss of lean mass. Whether this is deleterious remains unclear, but this change may have contributed to the unanticipated decline in inverted cling performance by the aged mice. In aged females, the Low Ile diet restored cardiac stroke volume and heart rate to levels comparable to Young Control-fed mice, and reversed the effects of age on phosphatidylglycerol levels; the overall effect on cardiac health is unclear, but these findings are consistent with emerging data suggesting that high blood levels of BCAAs are deleterious for cardiac function (Chen et al., 2019; Latimer et al., 2021; Portero et al., 2022; Uddin et al., 2019). While there was no significant effect of diet on cardiac output in males, the Low Ile diet significantly increased LV weight, and decrease mean and peak aortic flow velocity, which suggested the possibility of negative effects on cardiac function; this will need to be examined in detail in future studies.

The data we collected suggests that, particularly in males, a Low Ile diet may help prevent or slow age-associated increase in frailty in aging mice, much as we recently reported when this intervention was begun at 6 months of age (Green *et al*., 2023). While we were able to observe an age-dependent decrease in Barnes Maze performance, restriction of isoleucine or all amino acids did not improve performance; indeed the trend in both sexes was towards a decrease in performance with the Low Ile diet. This was at odds with our expectations, as we and others have found that PR and restriction of BCAAs improves cognition in a mouse model of AD, and we observed no negative effects of isoleucine restriction on cognition when begun at 6 months of age (Babygirija et al., 2023; Green *et al*., 2023; Tournissac et al., 2018). Larger cohorts studied for a longer period will be required to reach definitive answers on the impact of these diets on frailty and memory.

Low Ile and Low AA diets are highly effective at reverting age-associated changes at the molecular level. We found that both diets ameliorated age-related changes in lipid species in the female heart. In the liver, we observed that consumption of either a Low Ile or a Low AA diet had similar rejuvenating effects on multiple molecular aging rate indicators as observed in several other *bona fide* lifespan-extending treatments and genetic models. However, it is striking that a Low Ile diet was able to completely reverse the effect of aging on the molecular markers associated with the MAPK ERK and the CIT pathways, while a Low AA diet was not nearly as robust. This is provocative as a Low AA diet is also low in isoleucine, indicating the balance of dietary amino acids may play an important role in the regulation of aging. These differences may reflect in our recent findings that a Low Ile diet – but not a Low AA diet – extends lifespan when begun in young UM-HET3 mice (Green *et al*., 2023). Finally, taking an unbiased global approach, we find that most age-related changes in the hepatic transcriptome are reversed by a Low Ile diet in both sexes. Overall, the data presented here suggests that at the molecular level, a Low Ile diet has beneficial, perhaps rejuvenating, effects in aged mice.

There are a number of limitations in our study. First, as we sacrificed the animals for molecular analysis, we did not examine the effects of these diets on healthspan parameters at greater ages or on lifespan. The ultimate effect of these diets as late-life interventions on longevity will need to be determined in the future. While the molecular effects of these diets strongly suggest lifespan will be extended, and inhibition of mTOR signaling, which we observed here, starting late in life extends lifespan (Arriola Apelo et al., 2016; Harrison et al., 2009), we previously found that restriction of BCAAs starting in midlife did not extend lifespan (Richardson *et al*., 2021). In addition, the benefits of these diets to longitudinal phenotypes like frailty and cancer risk will benefit from additional studies with larger group sizes to come to firm conclusions on the impact of these diets on strength and biological robustness. Restriction of amino acids or isoleucine could potentially exacerbate sarcopenia, and thus this will also need to be carefully examined. The diets here strongly induce FGF21, which extends lifespan but is also associated with lower muscle mass (Roh et al., 2021; Zhang et al., 2012).

In summary, we have shown that restriction of all dietary amino acids or specifically restricting dietary isoleucine alone, interventions that have been demonstrated to extend both lifespan and healthspan when started in young and adult mice, can improve body composition and glycemic control in aged animals. The effects of these interventions on other aspects of healthspan are more mixed, tending to suggest a reduction in frailty but potentially neutral to harmful effects on muscle strength and cognition, which will require additional work to fully explore. Finally, we show that at the molecular level isoleucine restriction seems to rejuvenate the liver, phenocopying the molecular effects of several other interventions that extend lifespan while reversing age-related changes. Our results demonstrate that dietary composition – and in particular, the precise amino acids profile – is a critical regulator of healthy aging in aged mice. Overall, these results suggest that it may never be too late to obtain benefits on healthy aging from switching to a low protein or isoleucine restricted diet.

## ACKNOWLEDGEMENTS

We would like to thank Jiexian Chen and Josh Berg at the University of Michigan for assistance with molecular analysis of tissues, as well as all members of the Lamming lab for their assistance and input. The Lamming laboratory is supported in part by the NIH/NIA (AG056771, AG062328, AG081482, and AG084156 to D.W.L.), NIH/NIDDK (DK125859 to D.W.L.) and startup funds from the University of Wisconsin-Madison School of Medicine and Public Health and Department of Medicine to D.W.L. C-Y.Y. was supported in part by a training grant (T32AG000213) and a NIA F32 postdoctoral fellowship (F32AG077916). M.F.C. is supported by F31AG082504. R.B. is supported by F31AG081115. M.E.T. is supported by F99AG083290. C.L.G. is supported by HF AGE-009 from the Hevolution Foundation. MMS was supported in part by a Supplement to Promote Diversity in Health-Related Research RF1AG056771-06S1. HHP was supported in part by F31AG066311. JS is supported by the NIDDK (R01DK133479), JDRF (JDRF201309442 to JS), and JD is supported by an NSF GRFP. JS is a HHMI Freeman Hrabowski Scholar and is an American Federation for Aging Research grant recipient (A22068). RAM and GGG are supported by the Glenn Foundation for Medical Research. Support for this research was provided by the University of Wisconsin – Madison Office of the Vice Chancellor for Research and Graduate Education with funding from the Wisconsin Alumni Research Foundation. The authors used the UW-Madison Biotechnology Center Gene Expression Center (RRID:SCR_017757). The Lamming lab was supported in part by the U.S. Department of Veterans Affairs (I01-BX004031 and IS1-BX005524), and this work was supported using facilities and resources from the William S. Middleton Memorial Veterans Hospital. The content is solely the responsibility of the authors and does not necessarily represent the official views of the NIH. This work does not represent the views of the Department of Veterans Affairs or the United States Government.

## AUTHOR CONTRIBUTIONS

C-YY, MSG, TAH, RAM, and DWL conceived and designed the experiments. C-YY, LCSC, MSG, ITF, MFC, GGG, and ACR performed the experiments. C-YY, MSG, TAH, JWA, GGG, RAM, and DWL analyzed the data and wrote the manuscript.

## Materials & Methods

### Animals

All procedures were performed in accordance to institutional guidelines and were approved by the Institutional Animal Care and Use Committee of William S. Middleton Memorial Veterans Hospital (Madison, WI, USA) and the University of Wisconsin-Madison (Madison, WI, USA). C57BL/6J.NIA mice from the NIA Aged Rodent Colony were obtained at 20 months of age and at the young adult age of 6 months of age. Animals were provided *ad libitum* access to Laboratory Rodent Diet 5001 diet for 1-2 weeks and housed as 2-3 animals per cage before beginning their respective dietary interventions. All mice were maintained at a temperature of approximately 22⁰C, and health checks were performed daily by the facility staff. Mice were housed in a SPF facility in static microisolator cages under 12:12 light cycle conditions with ad libitum access to food and water unless specified below for upcoming experiments.

At the experiment start, aged animals were randomized at the cage level to groups of approximately equivalent weight, body composition and average frailty scores to one of three diets groups: Control with 21% calories derived from amino acids (TD.140711; Envigo), Low Ile with 67% less isoleucine content than the Control diet (TD.160734), and a Low AA diets with 7% calories derived from amino acids (TD.140714). The three diets are isocaloric, with supplemental non-essential amino acids used to balance the protein content in the Low Ile diet, and supplemental carbohydrates used to balance the calories content in the Low AA diet. The full composition of these diets is provided in **Table 1**.

Metabolic chambers indirect calorimetry was carried out using Oxymax/CLAMs metabolic chamber system (Columbus Instruments) for ∼48 continuous hours. The first ∼24 hours of data was discarded as acclimation period with a subsequent continuous 24 hours period utilized for data analysis. Food consumption monitoring was carried out in home cages over 2-4 days and analyzed using total animal mass in the cage. Body composition was determined using an EchoMRI Body Composition Analyzer.

### *In vivo* procedures

Glucose and insulin tolerance tests were performed by fasting the mice overnight for 16 hours or 4 hours in the morning respectively. In respective tests, mice were injected with glucose (1 g/kg; Sigma, G7021) or insulin (0.75 U/kg; Novolin) intraperitoneally (i.p.). Blood glucose was monitored via a Bayer Contour glucometer and test strips (Bayer, Leverkusen, Germany). All behavior tracking is done automatically with EthoVision XT program (Noldus). All following behavior/fitness testing follow previously described protocols (Bellantuono *et al*., 2020).

#### Novel Object Recognition

Briefly, a habituation/open field test is followed by the acquisition trial. Short-term memory test (STM) is carried out 1 hr after the acquisition trial, and the long-term memory test is carried out 23 hr from the acquisition trial. The habituation, acquisition, short-, and long-term memory test are 5 min each.

#### Barnes Maze Test

Briefly, the test is carried out on a maze table with 20 possible exit holes with distinct visible landmarks outside of the arena. Each animal is exposed to four acquisition days with a maximum trial time of 180 seconds. On day 5, a test trial takes place for short-term memory, then the mice are not experimented with until day 12, when a long-term memory test trial takes place.

#### Rotarod

Animals are trained in slow moving 4 rpm. On test trials, the accelerating rotarod (Rotamex 5, Columbus Instruments) is set to increase by 0.5 rpm per 4 seconds. There is a minimum rest period of 30 min between training and trials for each animal. The average trial performance for 3 test trials is taken for statistical analysis.

#### Inverted cling

Animals are placed onto a wire-grid bounded with masking tape and inverted onto the test bin. Mice naïve to the experiment are first trained through 3 trials of 30 seconds gripping prior to test trials. A minimum 30 min rest between training and, for each animal, 3 test trials are averaged for analysis.

#### Echocardiogram

Mice used for echocardiography were separate from the main study at a later age as indicated. Transthoracic echocardiography was performed using a Visual Sonics Vevo 770 ultrasonograph with a 30-MHz transducer as detailed previously (Harris et al., 2002). For acquisition of two-dimensional guided M-mode images at the tips of papillary muscles and Doppler studies, mice were sedated with 1% isoflurane administered through a facemask, hair removed, and maintained on a heated platform. Blood velocities across the mitral, aortic and pulmonary valves were measured using Doppler pulsed-wave imaging, angling the probe to obtain a nearly parallel orientation to the blood flow. End diastolic and systolic left ventricular (LV) diameter, as well as anterior and posterior wall (AW and PW, respectively) thickness were measured on line from M-mode images obtained in a parasternal long-axis view using the leading-edge-to-leading-edge convention. All parameters were measured over at least three consecutive cardiac cycles and averaged. Left ventricular FS was calculated as: ((LV diameterdias − LV diametersys)/LV diameterdias) × 100; ejection fraction: ((7.0/(2.4 + LV diameterdias)(LV diameterdias)3 − (7.0/(2.4 + LV diametersys) (LV diametersys)3/(7.0/(2.4 + LV diameterdias) (LV diameterdias)3 × 100; and LV mass: (1.05 × ((PWdias + AWdias + LV diameterdias)3 − (LV diameterdias)3)). Heart rate was determined from at least three consecutive intervals from the pulsed-wave Doppler tracings of the LV outflow tract. Ejection time was measured from the same outflow track tracings from the onset of flow to the end of flow. Isovolumic relaxation time was measured as the time from the closing of the aortic valve to the opening of the mitral valve from pulsed-wave Doppler tracings of the LV outflow tract and mitral inflow region. The same investigator obtained all images and measures.

#### FGF21 ELISA

Circulating FGF21 is measured using blood plasma obtained after 16 hr overnight fast and 3 hr refeed at 4 months after diet start, age 24 months for aged mice and age 10 months for young adult mice. Circulating FGF21 was quantified using a mouse/rat FGF21 quantikine ELISA kit (MF2100; R&D Systems, Minneapolis, MN, USA).

#### Western blots

Mice were sacrificed after overnight fasting and then refeeding all mice for 3-4 hrs with their respective diets. All tissues were flash frozen in liquid nitrogen. Liver lysate was homogenized in RIPA buffer with EDTA-free Protease and Phosphotase Inhibitor Mini Tablet (Thermo Scientific, A32961). Each blot was normalized to its control sample or the Aged Control group. Antibody product codes are as follows. Cell Signaling Technology: ULK1 (8054), pULK-S757 (14202), S6K1 (2708), pS6K1-T389 (9234), S6 (2217), pS6-S240/244 (2215), pS6-S235/236 (2211), eif2α (5324), peif2a-S51 (3597), 4E-BP1 (9644), and p4E-BP1-T37/46 (2855), AKT (9272), pAKT-T308 (2965), MK2 (12155), ACACA1 (4190) pAKT-S473 (9271), NDRG1 (9408), pNDRG1-T346 (5482), eIF4E (9742), peIF4E-S209 (9741), HSP70 (4872), MEK1 (12671), pMEK1-S217 (9154), ERK 1-2 (4695), pERK 1-2-T202 (9101), MNK1 (2195). Bioss: MNK2 (17697). Origene: TFAM (720347). Invitrogen: pMNK (700242), pMNK-T197 (700242). Abcam: FASN (22759), ACLY (40793). ABclonal: PLIN1 (A4758). ThermoFisher: PLIN3 (PA1-46161).

#### Quantative real-time PCR

rt-qPCR was carried out according to protocols described previously (Calubag et al., 2022) using TRI Reagent according to the manufacturer’s protocol. The primers that were used are as follows. P21 Fwd GAGACTAAGGCAGAAGATGTAGAG, Rev GCAGACCAGCATGACAGAT; P16 Fwd TGAGCTTTGGT TCTGCCATT, Rev AGCTGTCGACTTCATGACAAG; Il-1a Fwd TGCAGTCCATAACCCATGATC, Rev ACAAACTTCTGCCTGACGAG; Il-1b Fwd AGCCATGGCAGAAGTACCTG, Rev TGAAGCCCTTGCTGTAGT GG; Mcp-1 Fwd GATCTCAGTGCAGAGGCTCG, Rev TTTGCTTGTCCAGGTGGTCC; Il-10 Fwd ATAACTGCACCCACTTCCCA, Rev GGGCATCACTTCTACCAGGT; TNF-α Fwd ATGAGAAGTTCCCAAATG GC, Rev CTCCACTTGGTGGTTTGCTA; Il-6 Fwd CTGGGAAATCGTGGAAT, Rev CCAGTTTGGTAGCATC CATC;

#### Heart Lipidomics

MTBE extraction was performed with standards from SPLASH Lipidomix from Avanti, using methods outlined previously (Jain et al., 2022). Signal from the processed blank will be subtracted as background during data processing. For positive mode, samples were diluted to 15X in MeOH and 3uL were injected. Samples were run undiluted in negative mode and 5uL were injected. MS/MS data from pooled samples was run through Agilent LipidAnnotator, which were then used to processes MS1 data with Profinder (v8.0). Data was normalized to the internal standards and tissue weight, and further processed using R.

### Statistical Analyses

All statistical analyses were conducted using Prism, version 9 (GraphPad Software Inc., San Diego, CA, USA). Tests involving multiple factors were analyzed by either a two-way analysis of variance (ANOVA) with Time and Group as categorical variables or by one-way ANOVA with Group as the categorical variable followed by a Dunnett’s *post hoc* test for multiple comparisons against the Aged Control. Data distribution was assumed to be normal but was not formally tested.

### Transcriptomic Analysis

RNA was extracted from the liver using the PureLink RNA mini kit (Invitrogen, 12183025) with DNase (Invitrogen, 12185010) following manufacturer’s instructions. The concentration and purity of RNA was determined using a NanoDrop 2000c spectrophotometer (Thermo Fisher Scientific, Waltham, MA) and RNA was diluted to 100-400 ng/mL for sequencing. The RNA was then submitted to the University of Wisconsin-Madison Biotechnology Center Gene Expression Center & DNA Sequencing Facility, and RNA quality was assayed using an Agilent RNA NanoChip. RNA libraries were prepared using the TruSeq Stranded Total RNA Sample Preparation protocol (Illumina, San Diego, CA) with 250ng of mRNA, and cleanup was done using RNA Clean beads. Reads were aligned to the mouse (*Mus musculus*) with genome-build GRCm38.p5 of accession NCBI:GCA_000001635.7 and expected counts were generated with ensembl gene IDs (Zerbino et al., 2017).

Analysis of significantly differentially expressed genes (DEGs) was completed in R version 4.2.285 using edgeR (Robinson et al., 2010) and limma (Ritchie et al., 2015). Gene names were converted to gene symbol and Entrez ID formats using the mygene package. Male and female mice were analyzed separately, and one female outliers were removed following PCA analysis of the raw data. To reduce the impact of external factors not of biological interest that may affect expression, data was normalized to ensure the expression distributions of each sample are within a similar range. We normalized using the trimmed mean of M-values (TMM), which scales to library size. Heteroscedasticity was accounted for using the voom function, DEGs were identified using an empirical Bayes moderated linear model, and log coefficients and Benjamini-Hochberg (BH) adjusted p values were generated for each comparison of interest (α = 0.10). DEGs were used to identify enriched pathways, both Gene Ontology and KEGG enriched pathways were determined for each contrast, enriched significantly differentially expressed genes (FDR cutoff = 0.1, α = 0.05). All Gene Ontology categories (Cellular component, biological process, and molecular function) are presented together.

---

**Supplemental Figure 1.**
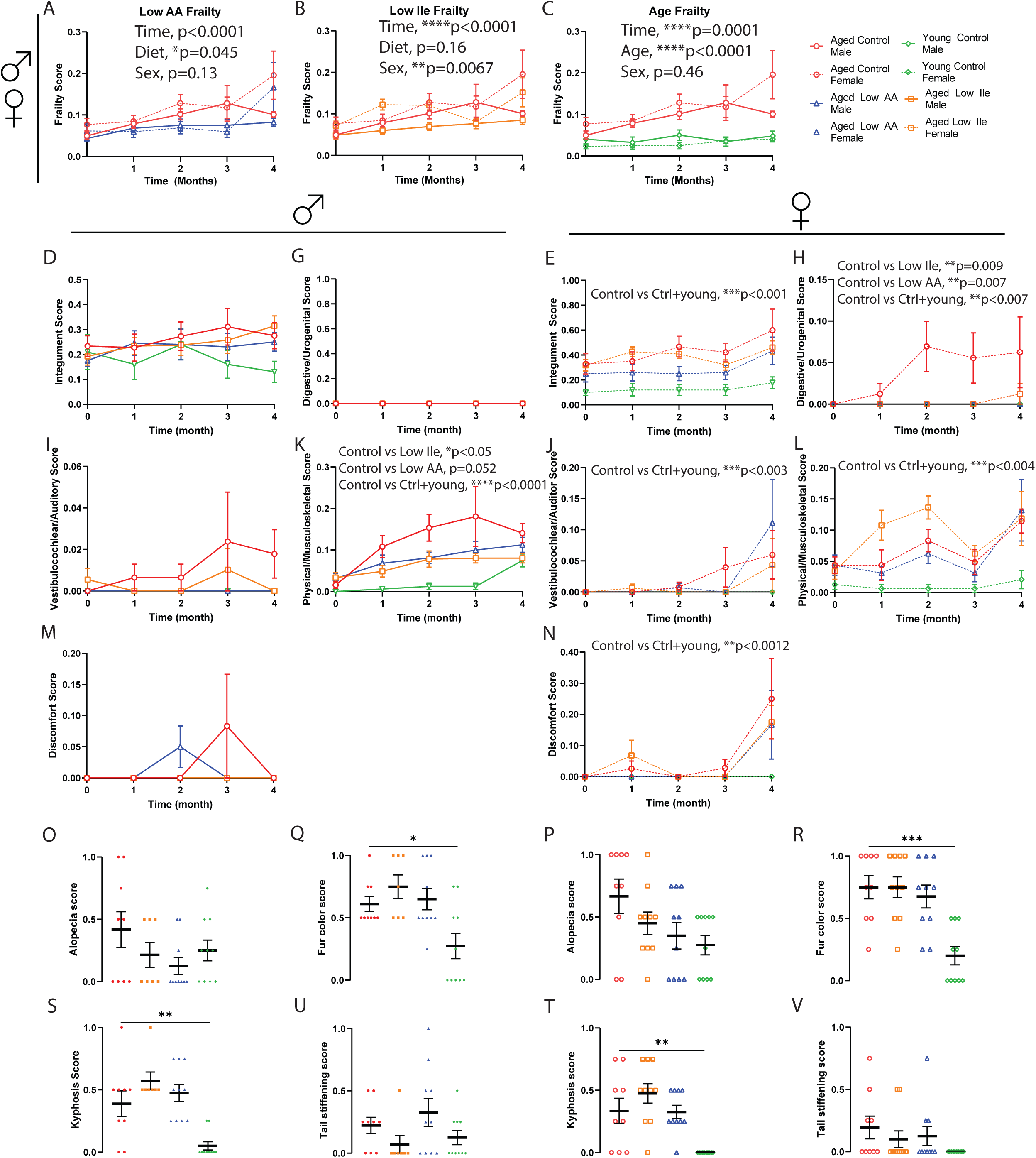
Frailty data 3-way ANOVA analysis and subcategories. **(A-C)** Three-way mixed-effects analysis of the frailty data as separated by the indicated factors. At the beginning of the experiments n=10-13/group; p-values represent the overall effect of time, diet, and sex. **(D-N)** Subcategory averages of the frailty data. n=10-13/group; p-values represent result of the 2-way mixed-effects analysis. **(O-V)** Selected individual frailty categories, presented as the average of 3- and 4-month scores. At the beginning of the experiments n=10-13/group, *p<0.05, **p<0.01, ***p<0.001, ****p<0.0001, ANOVA followed by Dunnett’s test. Data presented as mean ± SEM.

**Supplemental Figure 2.**
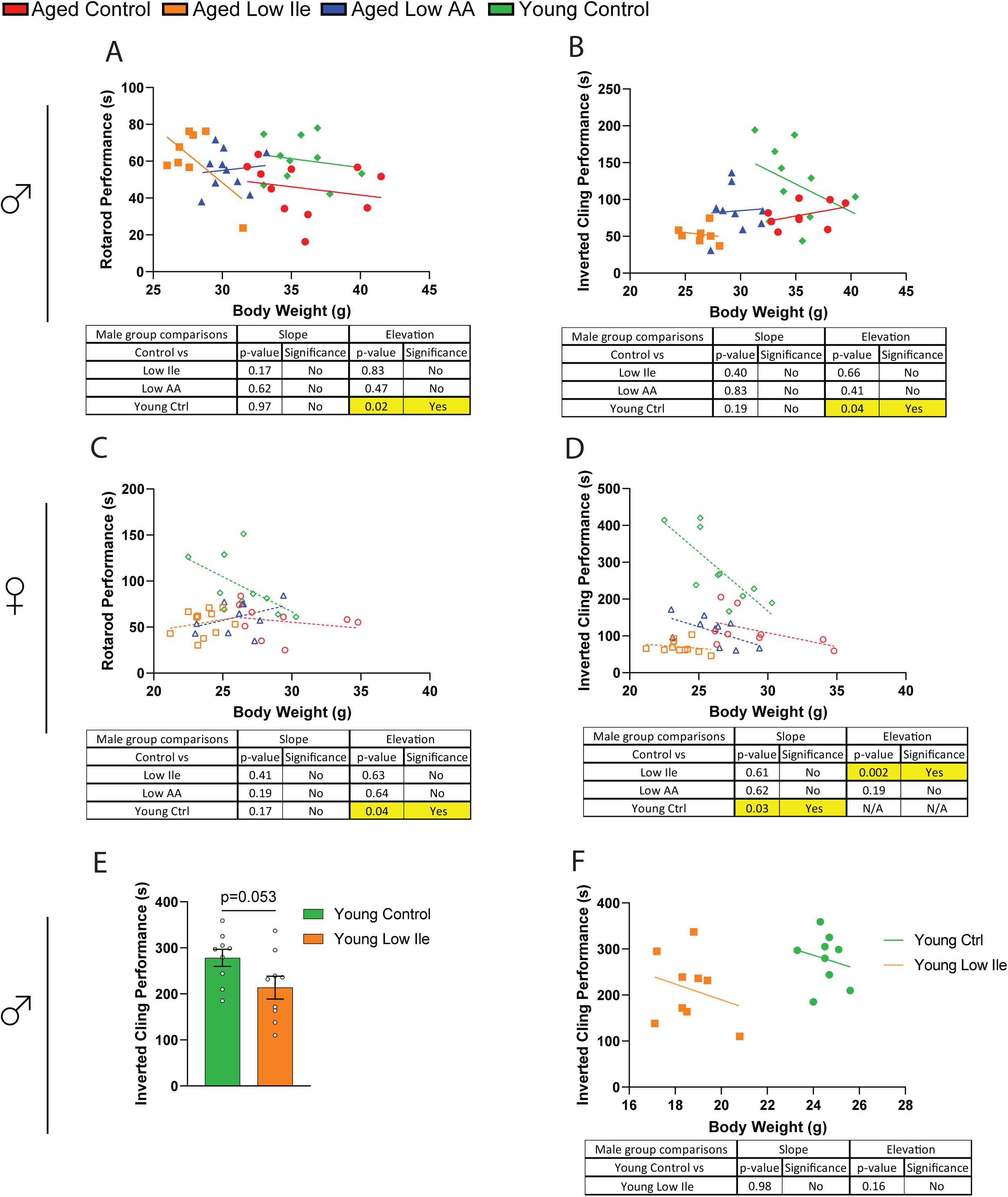
ANCOVA analysis of rotarod and inverted cling assay performance. **(A-D)** Rotarod and inverted cling performance as a function of body weight (n=8-11/group, slopes and intercepts were calculated using ANCOVA). **(E-F)** Young 3-month old male mice were fed either a Control or a Low Ile diet for at least 1 month before inverted cling assay (E), and inverted cling performance as a function of body weight (n=9/group, *p<0.05, t-test (E); slopes and intercepts were calculated using ANCOVA (F)). Data presented as mean ± SEM.

**Supplemental Figure 3.**
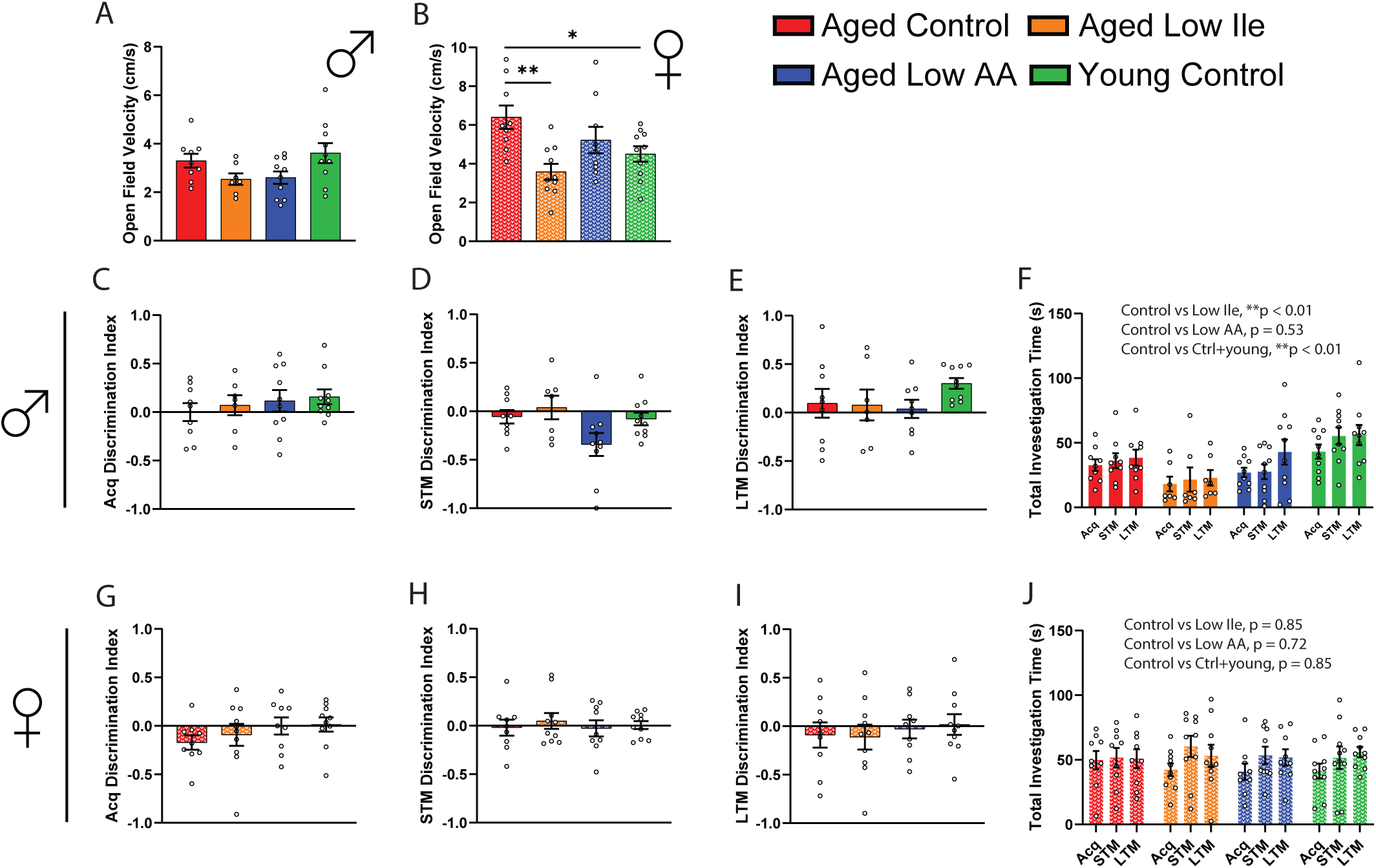
Open field and novel object recognition test. **(A-B)** Male (A) and female (B) mice in open field test. **(C-F)** Male mice novel object recognition test discrimination index in the acquisition phase (C), the short-term memory test (D), and the long-term memory test (E). (F) total investigation time in each trial. **(G-J)** Female mice novel object recognition test discrimination index in the acquisition phase (G), the short-term memory test (H), and the long-term memory test (I). (J) total investigation time in each trial. (A-E, G-I) n=7-10/group, *p<0.05, **p<0.01, ANOVA followed by Dunnett’s test. (F, J) n=7-10/group, p-values represent the main effect of diet from the indicated 2-way ANOVA. Data represented as mean ± SEM.

**Supplemental Figure 4.**
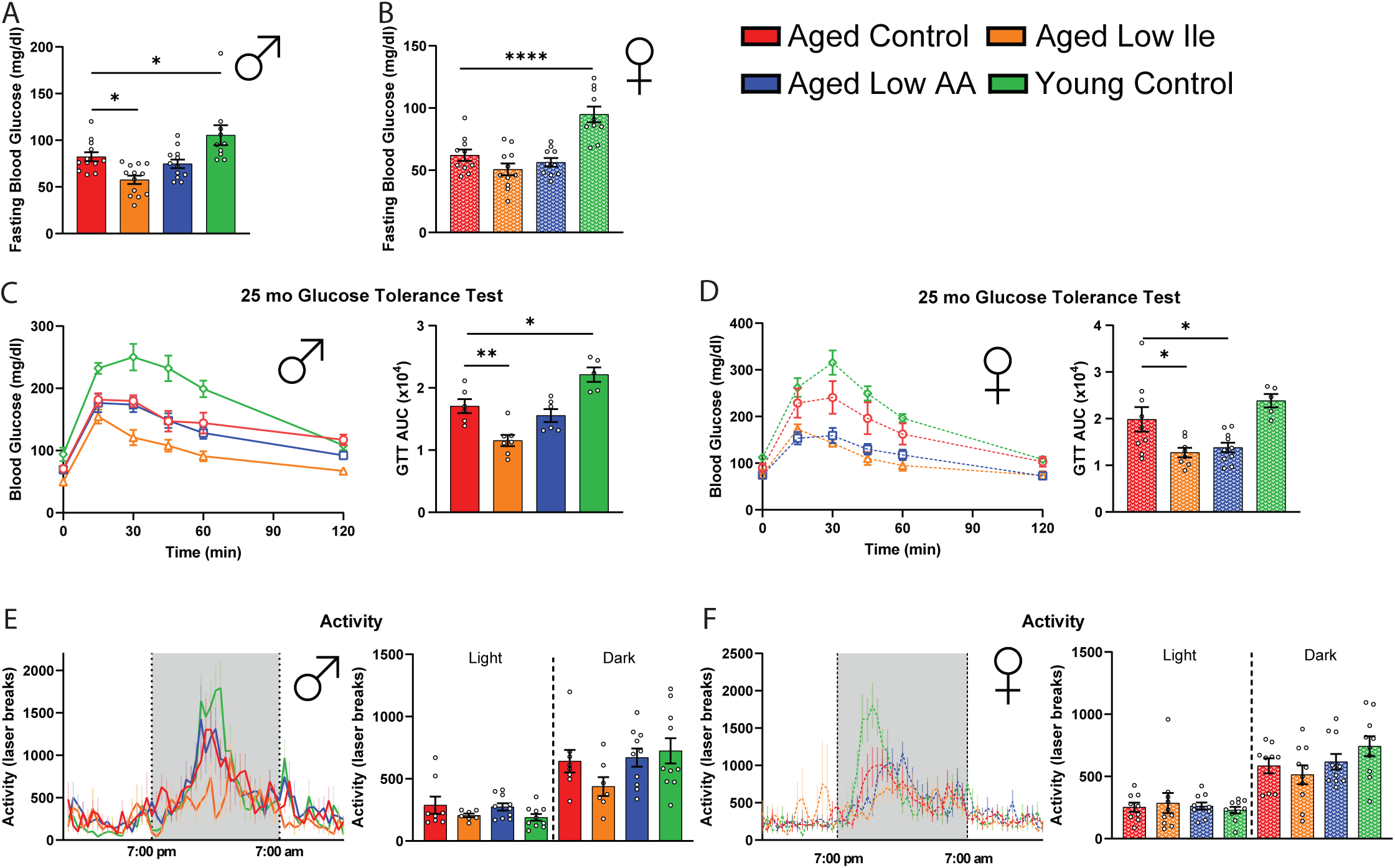
Effects of late-life Low AA and Low Ile diets on glycemic control and activity. **(A-B)** Fasting blood glucose of 21-month-old male (A) and female (B) mice after 3 weeks on the indicated diets. n=10-13/group. **(C-D)** Glucose tolerance of 25-month old male (C) and female (D) mice after 3 weeks on the indicated diets. n=5-10/group. **(E-F)** Spontaneous activity of male (E) and female (F) mice during the metabolic chambers experiments shown in Fig. 3. n=7-10/group. (A-F) *p<0.05, **p<0.01, ****p<0.0001, ANOVA followed by Dunnett’s test. Data represented as mean ± SEM.

**Supplemental Figure 5.**
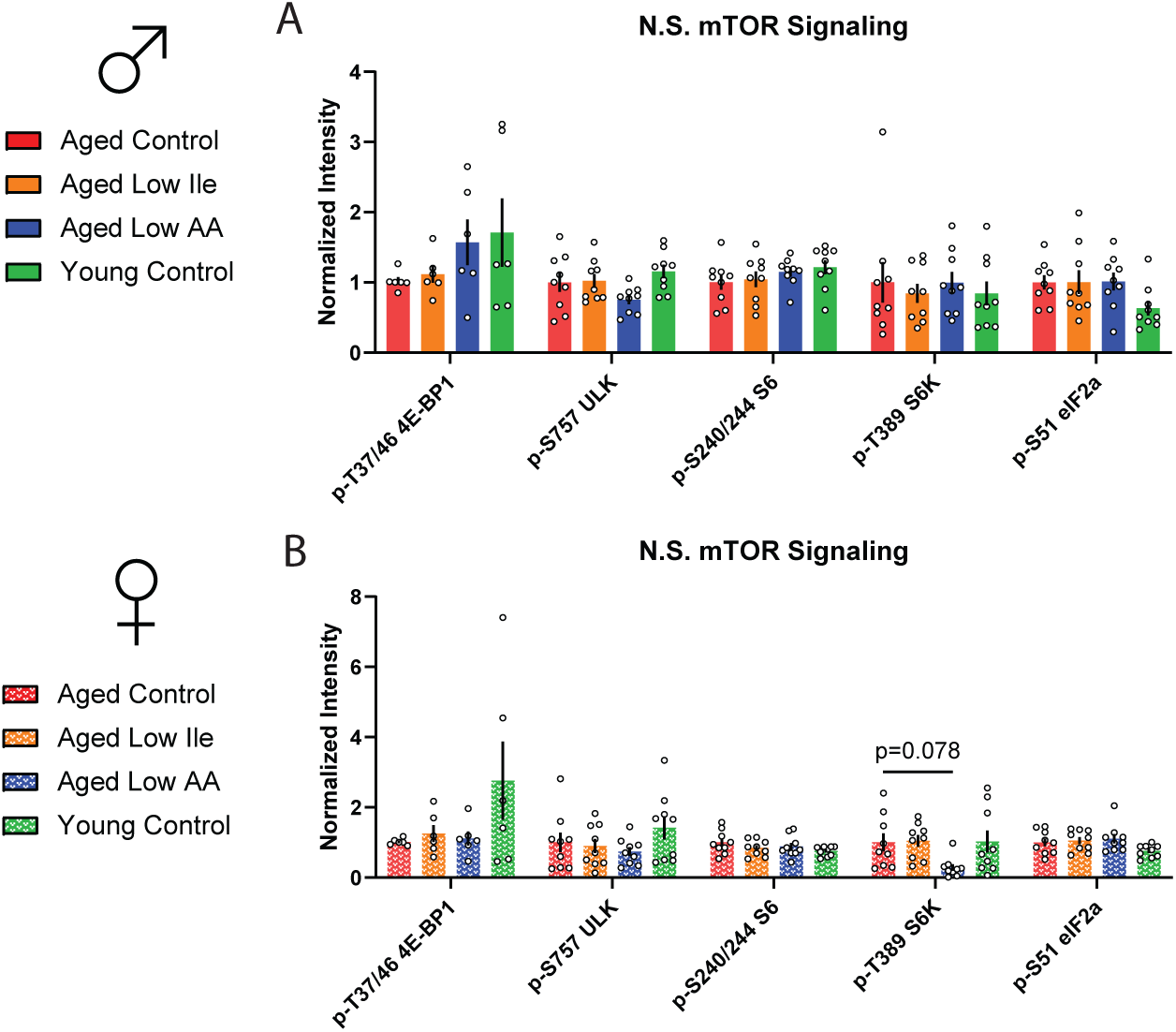
Non-significantly altered aging rate indicators in the aged mice liver. **(A)** Proteins not significantly altered by either diet or age in the livers of male mice. **(B)** Proteins not significantly altered by either diet or age in the livers of female mice. n=6-9/group, *p<0.05, ANOVA followed by Dunnett’s test. Data presented as mean ± SEM.

**Supplemental Figure 6.**
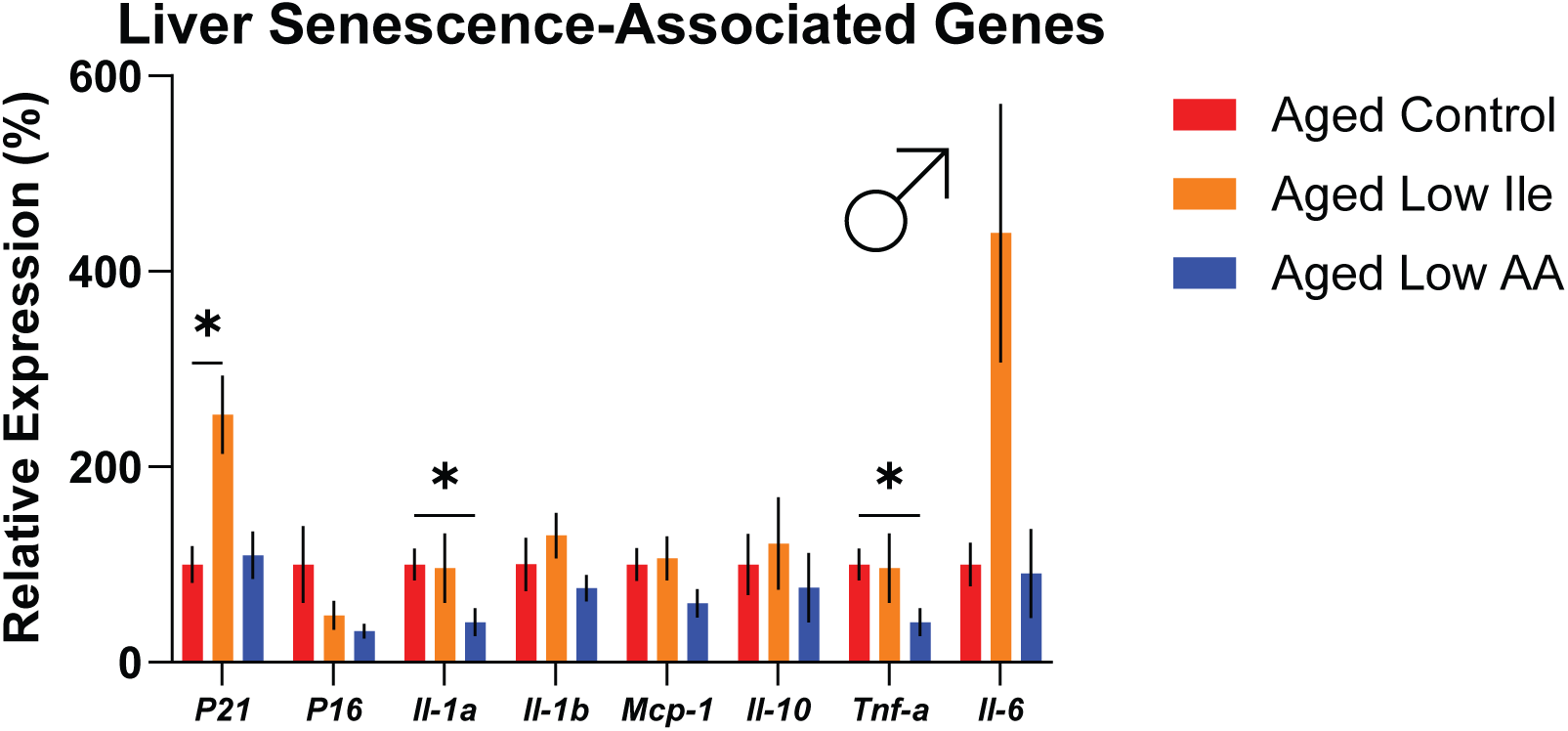
Expression analysis of senescence markers in the aged male liver. Expression of the indicated genes in the livers of 20-month-old mice on the indicated diets for 4 months was determined by qPCR. n=5-8/group, *p<0.05, ANOVA followed by Dunnett’s test. Data presented as mean ± SEM.

**Supplemental Figure 7.**
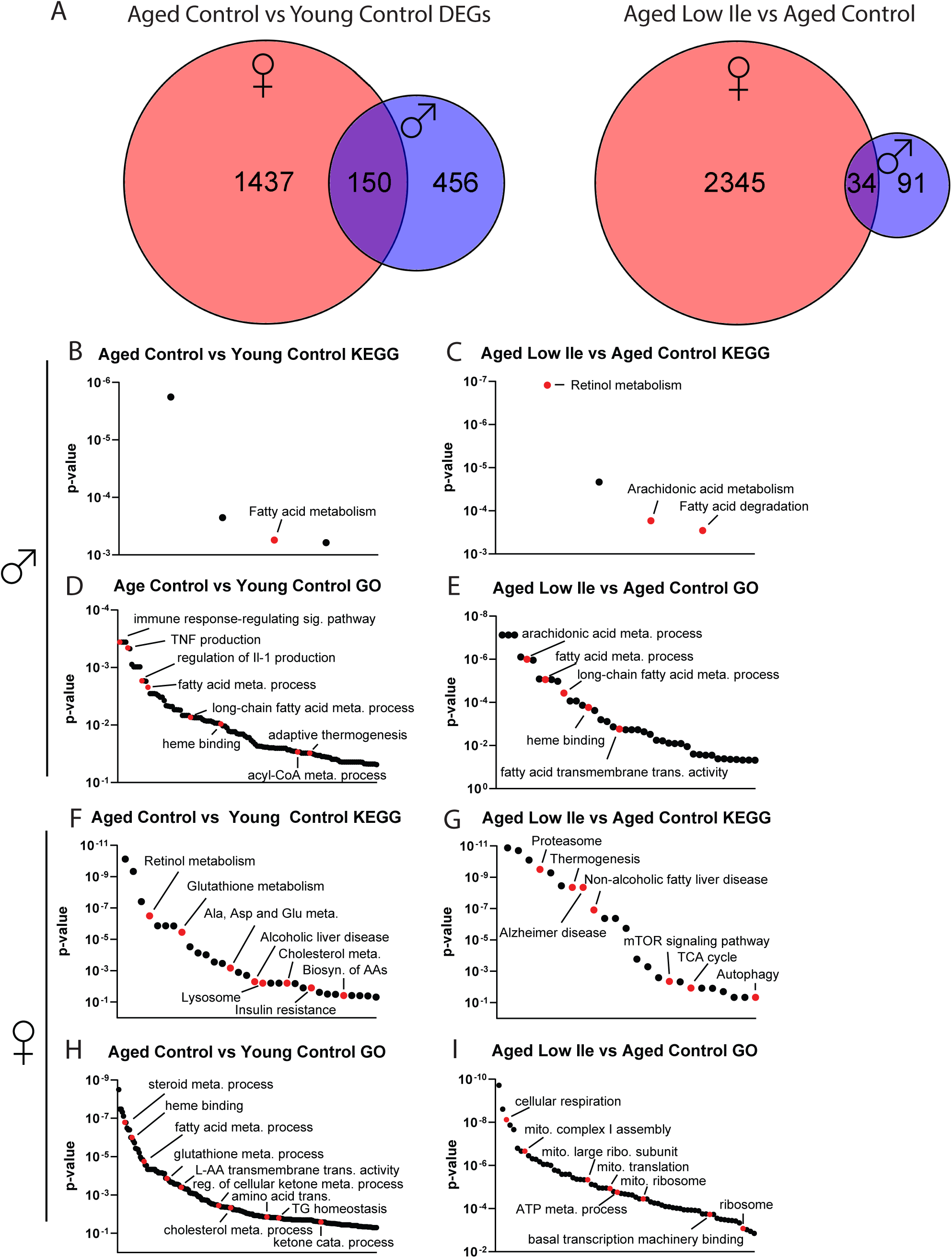
Venn diagram and enrichment analysis of differentially expressed hepatic genes. **(A)** Venn diagram showing the number of overlapping and non-overlapping DEGs between male and females. **(B-C)** Significantly enriched KEGG pathways by age (B) and diet (C) in male mice. **(D-E)** Significantly enriched GO terms by age (D) and diet (E) in male mice. **(F-G)** Significantly enriched KEGG pathways by age (F) and diet (G) in female mice. **(H-I)** Significantly enriched GO terms by age (H) and diet (I) in female mice. Transcriptomic analysis, n=5-6/group.

